# Design and Evaluation of Nanoscale Materials with Programmed Responsivity towards Epigenetic Enzymes

**DOI:** 10.1101/2024.03.26.585429

**Authors:** Priyanka Ray, Abbas Sedigh, Matthew Confeld, Lina Alhalhooly, Kweeni Iduoku, Gerardo M. Casanola-Martin, Hai Pham-The, Bakhtiyor Rasulev, Yongki Choi, Zhongyu Yang, Sanku Mallik, Mohiuddin Quadir

**Affiliations:** Department of Coatings and Polymeric Materials, North Dakota State University, Fargo, ND 58102, United States; Department of Pharmaceutical Sciences, North Dakota State University, Fargo, ND 58102, United States; Deapartment of Physics, North Dakota State University, Fargo, ND 58102; Department of Chemistry and Biochemistry, North Dakota State University, Fargo, ND 58102, United States; University of Science and Technology of Hanoi, Vietnam Academy of Science and Technology, 18 Hoang Quoc Viet, Cau Giay, Hanoi 10000, Vietnam

**Keywords:** Nanoparticles, stimuli-responsive nanoparticles, Histone deacetylase, enzyme-responsive nanoparticles, poly (acetyl L-lysine), computer-guided design, protein-ligand docking

## Abstract

Self-assembled materials capable of modulating their assembly properties in response to specific enzymes play a pivotal role in advancing ‘intelligent’ encapsulation platforms for biotechnological applications. Here, we introduce a previously unreported class of synthetic nanomaterials that programmatically interact with histone deacetylase (HDAC) as the triggering stimulus for disassembly. These nanomaterials consist of co-polypeptides comprising poly (acetyl L-lysine) and poly(ethylene glycol) blocks. Under neutral pH conditions, they self-assemble into particles. However, their stability is compromised upon exposure to HDACs, depending on enzyme concentration and exposure time. Our investigation, utilizing HDAC8 as the model enzyme, revealed that the primary mechanism behind disassembly involves a decrease in amphiphilicity within the block copolymer due to the deacetylation of lysine residues within the particles’ hydrophobic domains. To elucidate the response mechanism, we encapsulated a fluorescent dye within these nanoparticles. Upon incubation with HDAC, the nanoparticle structure collapsed, leading to controlled release of the dye over time. Notably, this release was not triggered by denatured HDAC8, other proteolytic enzymes like trypsin, or the co-presence of HDAC8 and its inhibitor. We further demonstrated the biocompatibility and cellular effects of these materials and conducted a comprehensive computational study to unveil the possible interaction mechanism between enzymes and particles. By drawing parallels to the mechanism of naturally occurring histone proteins, this research represents a pioneering step toward developing functional materials capable of harnessing the activity of epigenetic enzymes such as HDACs.

## 1. INTRODUCTION

Programmable nanomaterials capable of sensing and interacting with enzymes are generally composed of block copolymer assemblies, which recognize specific enzymes as destabilization triggers of their self-organized structures.^[1, 2]^ Such enzyme activities usually take place under mild aqueous or physiologically conducive conditions (aqueous, pH 5-8, 37 °C).^[3]^ Usually, the enzyme-nanomaterials interactions result in phase transition across different domains of the polymers forming the nanomaterials, leading to the gradual or catastrophic collapse of the assembled structure.^[4, 5]^ Enzyme-responsive nanomaterials has found useful applications in responsive soft materials design. ^[6-8]^ Specific application areas include biotechnology, agriculture, enzyme-catalysis, and medicine, where the materials can be used to form self-assembled platforms to encapsulate contents, such as small and macromolecular drugs,^[9, 10]^ diagnostic agents,^[3]^ and genetic materials^[11, 12]^. Mediated via programmed interaction with the destabilizing enzymes, these nanomaterials can control the exposure of the encapsulated content with their relevant targets, which can be of either biologic or non-biologic in origin. ^[13, 14]^ One of the biochemically important classes of epigenetic enzymes, which has not been harnessed earlier to produce enzyme-responsive nanomaterials is histone deacetylase (HDAC). These enzymes are found in nuclear and cytosolic fractions of cells, and are highly conserved across eukaryotic cells to carry out epigenetic processes, i.e., events that are manifested *via* the interactions between DNA and histone proteins.^[15-18]^ Mechanistically, HDACs remove an acetyl group from an *ε*-N-acetyl lysine amino acid on a histone protein, allowing the histones to wrap the DNA tightly.^[19, 20]^ We sought out to harness this properties of HDAC enzymes to design an enzyme-responsive nanomaterials, composed of block polypeptides, in the form of nanoparticles. We anticipate that, the use of HDAC as an activation trigger of a soft nanoparticles will open avenues to utilize and manipulate the expression of the HDACs - a critically important enzymes controlling cellular fate and disposition, in both plant and animal cells.^[21-23]^ This relevance stems from the fact that HDACs controls numerous epigenetic events associated with the evolution and progression of living cells.^[24]^ From therapeutic perspectives, the use of HDACs as a trigger for nanoparticles to release a therapeutic agent for rescuing or sensing genetically aberrant, diseased cells will, therefore, represent a paradigm shift in the scope of designing enzyme-sensitive therapeutic nanoparticles.^[21]^ To date, there is no report on HDAC enzymes being used as a destabilizing signal of nanoparticles. This is likely due to the difficulty in utilizing the unique function of HDAC to trigger nanoparticle disassembly (upon a deacetylation reaction). Therefore, in this work, we aim to set the design rules for nanoparticles that show conformational and morphological changes in the presence of HDACs by using HDAC8 as a representative member of this enzyme family.^[20]^ Furthermore, we show the potential of these nanoparticles in biomedical applications in terms of their safety, compatibility and efficacy. ^[25]^ Collectively, our study demonstrates for the first time the use of nanoparticles that relates structurally to naturally occurring histone proteins, which can interact with HDAC in a spatial and temporally-controlled pattern.

In addition to developing HDAC8-responsive block polypeptides, we also investigated the interactions of these polypeptides with HDAC8 enzymes computationally. Recently, computational, and molecular docking approaches have been routinely used in modern materials design workflow to help understand drug–receptor interactions. It has been shown in the literature that these computational techniques can strongly support and help the design of enzyme-sensitive materials by revealing the mechanism of molecular interactions between the substrate and the enzyme.^[26-32]^ Here, for proof-of-principle, we have used docking studies to study the binding orientations of poly(acetyl L-lysine), the nanoparticle-forming materials in our case, within the binding sites of HDAC8. The overarching scheme of this study is presented in **Figure 1**, illustrating a combined approach of experiment and computation via which we demonstrate that poly (ethylene glycol)-block-poly (acetyl L-lysine) block co-polypeptides can be used as building blocks to form the self-assembled soft materials with programmed sensitive to HDAC8 enzyme. The sensitivity of the polypeptide particles towards the enzyme was evidenced via the encapsulation and release of a reporter dye. We also demonstrated that the self-assembly of the polypeptides in the form of nanoparticles is mediated via the hydrophobic interactions taking place within the acetyl lysine-rich hydrophobic blocks, and HDAC8-mediated deacetylation of acetyl-L-lysine from the poly (acetyl L-lysine) block leads to the gradual loss of hydrophobicity of the block copolymer, destabilization of the nanoparticles, and release of the reporter content (**Figure 1A**). The purported chemical mechanism that drives the destabilization of HDAC8-sensitive nanoparticles is shown in **Figure 1B**. The usefulness and therapeutic compatibility of the HDAC-sensitive nanomaterials as a molecular transport platform was demonstrated using cancer-stem cells (CSCs) as representative and early in vitro models, where the growth and proliferation of these cells relies on HDAC enzymes and can be inhibited by Napabucasin, a STAT3 inhibitor^[33]^.

**Figure 1.**
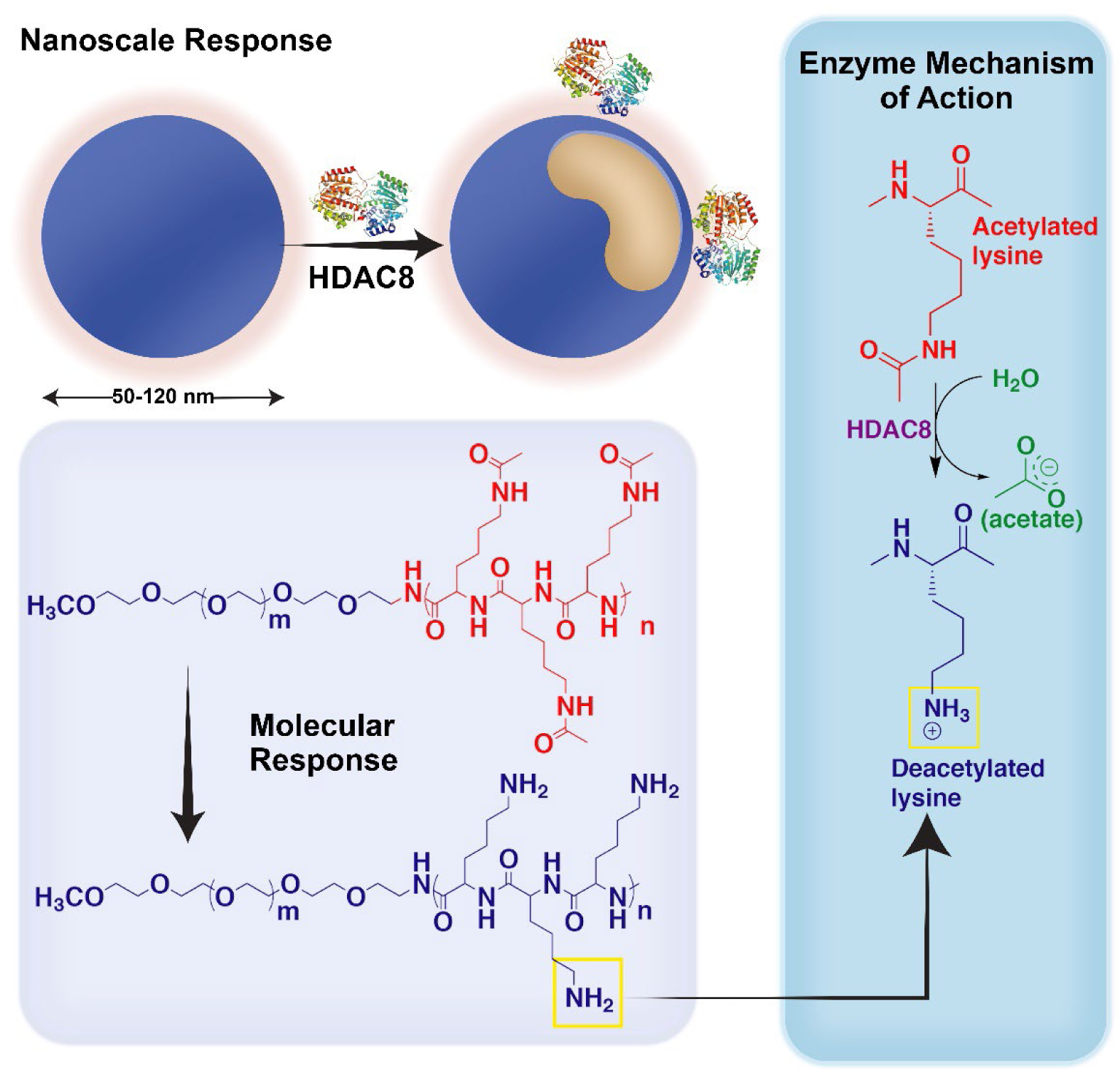
Schematic representation of HDAC-responsive nanoparticles and chemical architecture of nanoparticle-forming block copolypeptides. Mechanism of action of HDAC-responsive nanoparticles for content release under the influence of HDAC enzyme. (Left Panel): HDAC-sensitive nanoparticles are composed of block copolymer, PEG-block-poly (acetylated L-lysine), where the PEG constitutes the hydrophilic and poly (acetylated L-lysine) constitute the hydrophobic block; (Right Panel): The molecular mechanism of action of deacetylation of L-lysine by the HDAC enzyme of the hydrophobic block. Enzyme-mediated deacetylation is the primary driver of the amphiphilicity switch of the poly (acetylated L-lysine) block of the copolymer.

## 2. EXPERIMENTAL SECTION

### 2.1. Materials

Poly (ethylene glycol)-*block*-poly (L-lysine), henceforth abbreviated as PEG_m_-p (LysAc)_n,_ was purchased from Alamanda polymers (PEG = 5 kDa, m = 113 ethylene glycol units; poly (L-lysine, 33 kDa, n = 200 lysine residues, Structure, Figure 1A, structure 1). All other chemicals were purchased from Sigma-Aldrich, and anhydrous solvents were purchased from VWR EMD Millipore and were used without further purification. Fluor-de-Lys® fluorometric activity assay kit for detecting HDAC8 activity was obtained from Enzo Life Sciences. ^1^H NMR spectra were recorded using a Bruker 400 MHz spectrometer using TMS as the internal standard. Infrared spectra of synthesized compounds were recorded using an ATR diamond tip on a Thermo Scientific Nicolet 8700 FTIR instrument. Dynamic Light Scattering (DLS) measurements for determining the hydrodynamic diameter of nanoparticles were carried out using a Malvern instrument (Malvern ZS 90). UV-visible and fluorescence spectra were recorded using a Varian UV−vis spectrophotometer and a Horiba Fluoro-Log3 fluorescence spectrophotometer, respectively. TEM studies were carried out using a JEOL JEM2100 LaB6 transmission electron microscope (JEOL USA) with an accelerating voltage of 200 keV.

### 2.2. Synthesis and characterization of the HDAC8-responsive block polypeptides

HDAC8-responsive block polypeptide was synthesized *via* acetylation of **1**, *i.e.*, PEG_m_-p (LysAc)_n_ following the established protocol with minor variation.^[34]^ The molecular weight of the PEG block of the copolymer was 5 kDa (n = 113 ethylene glycol units), and the poly (L-lysine) block was 33 kDa (200 L-lysine residues). Briefly, 100 mg (0.0026 mmol) of the PEG-block-poly (L-lysine) was dissolved in a co-solvent composed of 4:1 DMF: 2,6-Lutidine (2,6-dimethylpyridine) solution (v/v). The solution was cooled in an ice bath, and 0.526 mmol acetic anhydride dissolved in DMF (1 mL) was added slowly. The reaction mixture was stirred at 0°C for 18 hours and precipitated into cold diethyl ether, followed by centrifugation at 8,000 rpm for 30 minutes two times. ^1^H-NMR and IR spectra were used to characterize the block co-polypeptide. We prepared three acetylated block copolymers from **1** at three degrees of acetylation, i.e., 25, 50, and 100% of all available lysine residues.

#### 2.2.1. Determination of the degree of acetylation

A ninhydrin test was performed on the starting material and the products to determine the number of lysine residues that have been acetylated. Typically, 2.5 mg of the polymers were dissolved in 2.5 mL of DMSO to which a few drops of freshly prepared ninhydrin solution (200 mg of ninhydrin in 10 mL ethanol) was added and heated in a water bath at 85°C till color development was observed. The absorbance (Abs) of each sample was measured, and the percentage functionalization was calculated using the following equation:

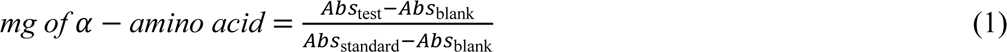

#### 2.2.2. Determination of the Critical Aggregation Concentration (CAC) of HDAC8-responsive block co-polypeptides

To evaluate the aggregate forming capacity of the amphiphilic polypeptides and to estimate the stability of these aggregates in an aqueous environment, we determined the critical aggregation concentration (CAC) of PEG-block-poly (L-acetylated Lysine) co-polypeptides. A stock solution of 0.1 mM pyrene in dichloromethane was prepared, and an aliquot of 10 μL of this solution was taken in a set of vials from which dichloromethane was allowed to evaporate by air-drying overnight. Various measured amounts of acetylated co-polypeptides were added to each of these vials (from a stock solution of 10 µM). The block co-polypeptide concentrations varied from 0.15 to 5.5 µM, with the final concentration of pyrene in each vial maintained at 1 μM. The vials were sonicated for 90 min and then allowed to stand for 3 h at room temperature (r.t.) before recording the fluorescence emission spectra at an excitation wavelength of 337 nm with slit widths of 2.5 nm (for both excitation and emission). The ratio of the intensities at 373 and 384 nm was plotted against the co-polypeptide concentration, and the curve’s inflection point was used to determine the CAC as per published procedures^[35, 36]^.

### 2.2.3. Preparation and characterization of HDAC8-responsive nanoparticles from PEG-block-poly (acetylated L-lysine co-polypeptides)

To prepare self-assembled structures from the co-polypeptides, we employed a non-solvent induced phase separation (otherwise known as nanoprecipitation or solvent shifting) method from a selective solvent (DMSO, for both blocks) to a non-selective solvent (buffer, to a single block).^[37-39]^ The acetylated co-polypeptides were dissolved in 250 μL of DMSO, and the solution was added dropwise to 750 μL of PBS buffer (pH 7.4). The resultant solution was transferred to a Float-a-Lyzer (MWCO 3.5−5 kDa) and dialyzed against 800 mL PBS buffer (pH 7.4) overnight with constant agitation at moderate speed. Dialysis over a stipulated period resulted in the formation of nanoparticles. Hydrodynamic diameters of resulting particles prepared from the acetylated block copolymers were determined using Dynamic Light Scattering (DLS) at a scattering angle of 90°. Surface charge or Zeta (ζ-) potential of block co-polypeptides was measured by evaluating the electrophoretic mobility of samples with a nanoparticle concentration of 10 mg/mL. An average of 5 readings were acquired to identify the zeta potential, and for all measurements, sample solutions were filtered through 0.45 μm PES filters.

### 2.2.4. Transmission electron microscopy (TEM) imaging of co-polypeptide nanoparticles

A drop of nanoparticle sample obtained from the self-assembly of PEG-block-poly (acetylated L-lysine) co-polypeptide was placed on a 300-mesh Formvar carbon-coated copper TEM grid (Electron Microscopy Sciences) for 1 min and wicked off. Phosphotungstic acid 0.1%, pH adjusted between 7.0−8.0, was dropped onto the grid, allowed to stand for 2 min, and then wicked off. Nanoparticles (untreated or treated with h HDAC8 enzymes) were imaged for their microstructure via TEM at 200 keV.

### 2.2.5. Encapsulation of 5(6)-Carboxyfluorescein (CF) in HDAC8-responsive Nanoparticles

5(6)-Carboxyfluorescein was encapsulated by the following procedure: 10 mg of the co-polypeptide and 1 mg of 5(6)-Carboxyfluorescein was dissolved in 250 µL DMSO, and the solution was then added dropwise to a 750 µL PBS buffer solution under magnetic stirring. This was left stirring for an hour at room temperature, followed by dialysis (MWCO 1−1.5 kDa) against 800 mL PBS buffer with regular media change till no further discoloration of the media was observed. 20 μL (1%) of Triton was added to disintegrate the polymersomes, and the fluorescence emission intensity was measured for total release after disintegration, considered as the initial intensity at *t* = 0.

### 2.3. Biochemical activity evaluations

#### 2.3.1. Preparation of HDAC8 enzymes and the assay buffer

HDAC8 was obtained from Professor D. K. Srivastava’s laboratory (Chemistry and Biochemistry, NDSU). The enzyme was prepared and purified according to the protocol described by Srivastava laboratory in earlier publications. Briefly, the cDNA containing plasmid (mammalian expression vector pCMV-SPORT) was purchased from Open Biosystems Huntsville (clone ID 5761745). The HDAC-8 gene was amplified by PCR reaction using forward and reverse primers. Following ligation of the PCR product with pLIC-His expression vector (obtained as a gift from Prof. Stephen P. Bottomley, Monash University, Australia), the recombinant plasmid (pLIC-His6-HDAC8) was transformed into *E. coli* BL21 codon plus DE3 (RIL) chemically competent cells (purchased from Stratagene^TM^ California) for expressing the HDAC8 enzyme. The transformed cells were cultured in LB medium at 37°C to reach OD600. At this point, the culture was supplemented by 100 µM ZnCl_2,_ and the culturing was continued at 16 °C for 16 additional hours. Cells were harvested by centrifugation at 5,000 rpm. The cell pellet was sonicated using lysis buffer, and the resulting lysate was centrifuged at 15,000 g for 30 min at 4°C. The suspension was filtered to remove cell derbies, and pure HDAC8 enzyme was obtained using the HisTrap column on AKTA purifier UPC 10 (GE Healthcare Life Sciences). SDS-PAGE agarose gel and catalytic activity analysis confirmed the presence of the HDAC8 enzyme. The HDAC8 assay buffer used for our studies had the following composition: 50 mM Tris-HCl buffer containing 137 mM NaCl, 2.7 mM KCl, 1 mM MgCl_2_, 1 mg/ml BSA, pH 7.5 while the HDAC8 lysis buffer comprised of 50 mM Tris-HCl, 150 mM KCl, 3 mM MgCl2, 1mM 2-mercaptoethanol, 1 mM PMSF (phenylmethylsulphonyl fluoride) and 0.25 % Triton X-100, pH 8. Formally, the HDAC8 enzyme activity was monitored in the assay buffer by Fluor-de-Lys (R-H-K(Ac)-K(Ac)-AMC) fluorogenic substrate (KI-104) *via* trypsin coupled assay established by Schultz and coworkers^[16]^ (Shown later in **Figure 4**).

#### 2.3.2. Fluor-de-Lys® HDAC fluorometric assay

The assay was performed on a microplate reader. Varying polymersome concentrations were used, and the HDAC8 concentration was optimized to be 850 nM. The Fluor-de-Lys® substrate concentration was maintained at 200 µM. The excitation wavelength was 360 nm, and the emission was 460 nm. The emission spectra were recorded for 4 hours.

#### 2.3.3. Release of 5(6)-Carboxyfluorescein in HDAC8-responsive Nanoparticles under the influence of the enzyme

The rate and extent of release of carboxyfluorescein from HDAC8-responsive nanoparticles were evaluated by tracking the fluorescence emission intensity of the dye at 518 nm with an excitation wavelength of 480 nm. In a representative experiment, HDAC8 enzyme was added to a polymersome suspension in HDAC8-assay buffer (please see earlier section) to achieve the final concentrations of HDAC8 as 50 nM, 100 nM, and 1µM. The fluorescence intensity was measured every 7 minutes for over an hour and later time points (8h) at 25°C. The experiment was conducted in triplicate, and the data were collected with a fluorescence spectrophotometer. The fractional release of carboxyfluorescein was calculated by comparing the fluorescence intensity of the nanoparticle solution to the intensity of a solution containing an equal concentration of nanoparticles in the presence of 1% Triton X (without HDAC enzyme). The concentration of Triton X required for the complete release of encapsulated CF from the nanoparticles was adjusted beforehand by confirming that increasing the reagent concentration did not cause a further increase in the fluorescence intensity of the CF. The fractional release of CF from the HDAC8-responsive nanoparticles was calculated as the ratio of the difference between the fluorescence intensity of carboxyfluorescein at time t (It) and its initial intensity at t = 0 (I0) and the difference between the intensity upon treatment with 1% Triton X-100 (Triton) and the intensity at t=0 according to an earlier published report^[40]^.

#### 2.3.4. Atomic force microscopy and nanoparticle tracking analysis of HDAC8-responsive nanoparticles

HDAC8-responsive nanoparticles (prepared from 10 mg/mL of block co-polypeptides) were first analyzed for the particle diameter in the absence and presence of HDAC8. The size distribution and concentration of nanoparticles were determined by nanoparticle tracking analysis (NTA) using the NanoSight NS300 system (Malvern Pan Analytical Ltd, UK). The exosome samples were diluted to 1000-fold in PBS for NTA measurements. The samples were infused with the syringe pump at a constant speed of 20 into the microfluidic flow cell equipped with a 532 nm laser and a high-sensitivity scientific CMOS camera. At least three videos per sample were recorded with a camera level of 11 - 13 for 30 s at 25°C. All data were analyzed using NTA software (version 3.4) with a detection threshold of 5. Atomic force microscopy (AFM) imaging of nanoparticles was performed using a commercial atomic force microscope (NT-MDT NTEGRA AFM). The samples were prepared by incubating 10 µl of nanoparticle solution on silicon substrates for 30 min in a sealed compartment to protect against evaporation at room temperature. The samples were then rinsed with de-ionized water (Millipore) and dried under N_2_ flow. The samples were imaged under ambient conditions in semi-contact mode with a resonant frequency of 190 kHz AFM probes (Budget sensors).

#### 2.3.5. Encapsulation of a therapeutically-relevant molecule in HDAC8-responsive nanoparticles as proof of concept

We used Napabucasin (NAPA) as the model drug to demonstrate the proof-of-concept of the utility of the co-polypeptide nanoparticles as a drug delivery platform. This is because NAPA affects the growth and proliferation of CSCs, which we found to have enriched HDAC8 activity (please see Figure 5). To prepare the NAPA-encapsulated system, 10 mg of the block co-polypeptide and 5 mg of NAPA were dissolved in 250 μL of DMSO. The solution was added dropwise to 750 μL of PBS (pH 7.4) with constant stirring. The solution was stirred overnight, followed by filtration using an ultracentrifuge filter (MWCO 3.5−5 kDa) at 5000 rpm for 3 h to prepare the purified nanoparticles. The resulting nanoparticle suspension was dispersed in chilled (4 °C) buffer to a concentration of 10 mg/mL, and the filtrate was used to quantify the amount of the drug. The encapsulation efficiency (EE %) was calculated using the following equation:

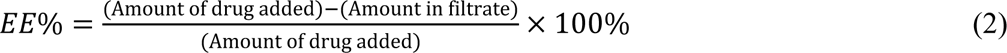

#### 2.3.6. In Vitro enzyme-mediated Drug Release

In vitro drug release was studied simultaneously in the absence and presence of 1µM HDAC8 and using the assay buffer. Napabucasin, a STAT3 inhibitor, was used as the model drug. An aliquot of 1 mL of the drug-encapsulated polymersomes was taken in one Float-a-Lyzer (MWCO 3.5−5 kDa) chamber and was dialyzed against 5 mL of HDAC assay media. After a specified time interval, 1 mL of sample was withdrawn and replaced with the same volume of the fresh assay media. The samples were then analyzed for NAPA concentration using UV−vis spectroscopy.

#### 2.3.9. Cell-culture and cytotoxicity assay

In-vitro studies were carried out using four cell variants. HPNE, PANC-1, and MIA PaCa-2 cell lines were obtained from American Type Tissue Culture (ATCC) and maintained and passed according to ATCC recommendations. The fourth cell variant is patient-derived xenograft pancreatic cancer stem cells (CSCs) obtained from Celprogen. The stem cells were maintained and passaged using Celprogen-recommended media, flasks, and plates. All studies used a passage number of less than 10 except for the stem cells, where only passages below 5 were used. Cells were seeded in a 96-well plate at a density of 10,000 cells/well and allowed 24 hours of incubation before adding treatments. Nanoparticles containing napabucasin were suspended in serum-free cell culture media and added to their respective wells. Twenty-four hours post-treatment, the cells were washed 3x with phosphate-buffered saline, and a 10% Alamar Blue (Bio-Rad) concentration was used to determine cell viability/cytotoxicity.

#### 2.3.10. Molecular docking computational studies of HDAC8 enzyme with the block polypeptide

To identify the putative interactions of HDAC8 responsive block copolymers and their nanoparticle ensemble with the target enzyme, HDAC8, we conducted computational molecular docking studies using ICM-pro 3.8.3 (Molsoft, L.L.C.).^[41]^ The coordinates for HDAC8 receptors were retrieved from Protein Data Bank (PDB ID: 2V5W and 1T64) ^[42, 43]^ and then pretreated following the standard procedures, including the exclusion of water molecules and heteroatoms, addition of hydrogen and charges using ECEPP/3 templates, correction of side chain and missing loop.^[44]^ As several water molecules could be part of the binding interaction network, 16 water molecules inside the pocket were kept for docking assays. Additionally, the configuration of the critical residues forming the HDAC8 active site tunnel, such as His142, His143, Gly151, Phe152, His180, Phe208, Pro273, and Tyr306, were checked and corrected.^[45]^ As 2V5W was reported with a mutation at position 306,^[42]^ we tried to induce a Phe306Tyr mutation and reoptimize the protein, 2V5W, taking 1T64 for structural alignment. This mutated target was then used for docking assays. For ligand preparation, a monomer structure of PEG_m_-p (LysAc)_n_ was generated and optimized by using the Molecular Editor wizard in ICM. The docking study was then performed, generating 1000 conformations. The top-scoring poses were ranked and selected based on the ICM scoring function, whose variables consist of the conventional interactions and internal force field energy, i.e., hydrophobic van der Waals, electrostatic interaction, solvation/desolvation, conformational loss energy, and hydrogen bonding.^[44]^

## 3. RESULTS AND DISCUSSION

This work’s central hypothesis is that nanoparticles composed of hydrophobic domains rich in acetylated L-lysine will act as a substrate for a deacetylating enzyme, such as histone deacetylase. The deacetylation reaction will lead to the reversal of solubility of the hydrophobic block and subsequent destabilization of nanoparticles. To prove this hypothesis, we designed a poly (ethylene glycol)-block-poly (acetylated L-lysine) block co-polypeptide abbreviated as PEG_m_-p (LysAc)_n_, where m and n represent the number of repeating units for the respective blocks. Within the HDAC family, we selected HDAC8 as a representative member of the deacetylating enzyme to prove our hypothesis. This enzyme is a member of the class I HDACs and is localized both in the nucleus and the cytosol, involved in numerous epigenetic and transcriptional processes related to health and disease.^[22]^ Due to the overexpression of HDAC8 with various pathophysiological conditions, HDAC8-responsive nanoparticles can potentially be used in drug delivery.^[46-48]^ As such, we also proved that HDAC8-responsive nanoparticles can be used as a drug delivery vehicle by encapsulating a STAT3 inhibitor, Napabucasin (NAPA), inside the particles.^[49]^ NAPA has been a drug of choice to suppress hard-to-kill cancer stem cells (CSCs).^[50-52]^ We envisioned that the destabilization of HDAC8-responsive nanoparticles by HDAC8 would induce NAPA (a STAT3 pathway inhibitor) release in a time-dependent pattern.^[53-55]^

### 3.1. Synthesis of block co-polypeptides with acetylated L-lysine side chain

Acetylated L-lysine is the substrate of the HDAC8 enzyme. Therefore, we acetylated the L-lysine residue of a PEG-block-poly (L-lysine) block copolymer. Acetylation of the poly (L-lysine) domain resulted in the formation of an amphiphilic block copolymer, which was later used to create self-assembled nanoparticles. PEG-conjugated and ε-amino group protected poly (L-lysine) are important block copolymer candidates for forming colloidal nanoparticles for drug and gene delivery applications and as functional materials for tissue engineering.^[36, 56-58]^ We prepared HDAC8-accessible, PEG-block-poly (acetylated L-lysine) block copolymer by reacting the pre-formed PEG-block (poly L-lysine) (M_n_ = 21 kDa, PEG = 5 kDa) with acetic anhydride in the presence of pyridine (**Scheme 1**).

**Scheme 1:**
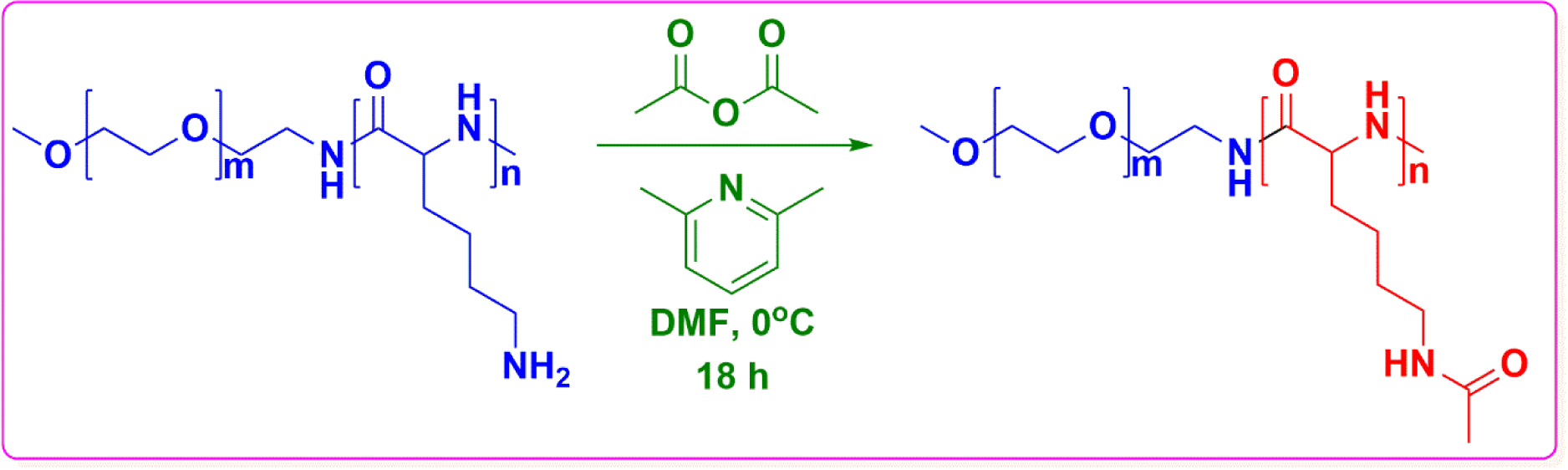
(A) Synthesis of the block copolymer, PEG-block-poly(acetylated L-lysine) composed of 113 units (m) of ethylene glycol and 200 L-lysine residues. Full functionalization results in a degree of acetylation of 86% calculated from ^1^H-NMR and Ninhydrin tests, indicating 172 L-lysine units (n) that can be acetylated by this method.

The method, initially described by Thoma et al., provided the advantage of more than 80% conversion of the available ε-amino groups of lysine residues present in the copolymer’s 16 kDa poly(L-lysine) block. Acetylated block copolymers were characterized, and the degree of acetylation was calculated via ^1^H NMR (Supporting information, Figure S1) and by Ninhydrin test (Supporting information, Figure S2). The latter test demonstrated that the degree of acetylation obtained under the reaction conditions specified in **Scheme 1** yielded a degree of acetylation of 86% for the PEG-block-poly(L-lysine) block copolymer bearing 113 ethylene glycol units (in the PEG block) and 200 acetylated L-lysine residues. This indicates that, of 200 available lysine residues, 172 residues were acetylated. Further, we evaluated the critical aggregation concentration (CAC) of the newly synthesized PEG-block-poly (acetylated L-lysine) block copolymer using fluorescence spectroscopy-based methods using pyrene as a probe. The CAC value is a robust indication of the stability of any nanoscale aggregates that amphiphilic copolymers usually form. The ratio of the first (λ = 373 nm) and third (λ = 384nm) peaks in the fluorescence emission spectra of pyrene is recorded in the presence of varying concentrations of the copolymer, and the ratio of I_373_/I_384_ is determined and plotted as a function of the copolymer concentration. We observed that, depending on the degree of acetylation, the intensity ratio decreased up to a particular concentration of the copolymer, after which it remained unchanged, independent of concentration (Figure S3, Supporting information). From this experiment, CAC values of PEG-block-poly (acetylated L-lysine) were found to be within the range of 2.2 × 10^−6^ M (for n = 200 L-lysine units bearing polymers, degree of acetylation = 86%). We also observed that block copolymers with reduced levels of acetylation showed higher CAC value (5.26 × 10^−6^ M for block copolymers with a degree of acetylation of 25%). This result indicated that acetyl side chains increased the stability of nanoscale aggregates of

PEG-block-poly (acetylated L-lysine) copolymer, most likely due to the increased capacity of these side chains to form H-bonding and hydrophobic interactions with each other.

### 3.2. Nanoscale features of HDAC8-responsive nanoparticles

The amphiphilicity of PEG-block-poly (acetylated L-lysine) drives the self-assembly of this copolymer to form nanoparticles under solvent exchange (nanoprecipitation) conditions. Using dynamic light scattering (DLS), we evaluated the particle size (in terms of hydrodynamic diameter) and surface charge (ζ-potential) of these nanoparticles (**Figure 2**).

**Figure 2.**
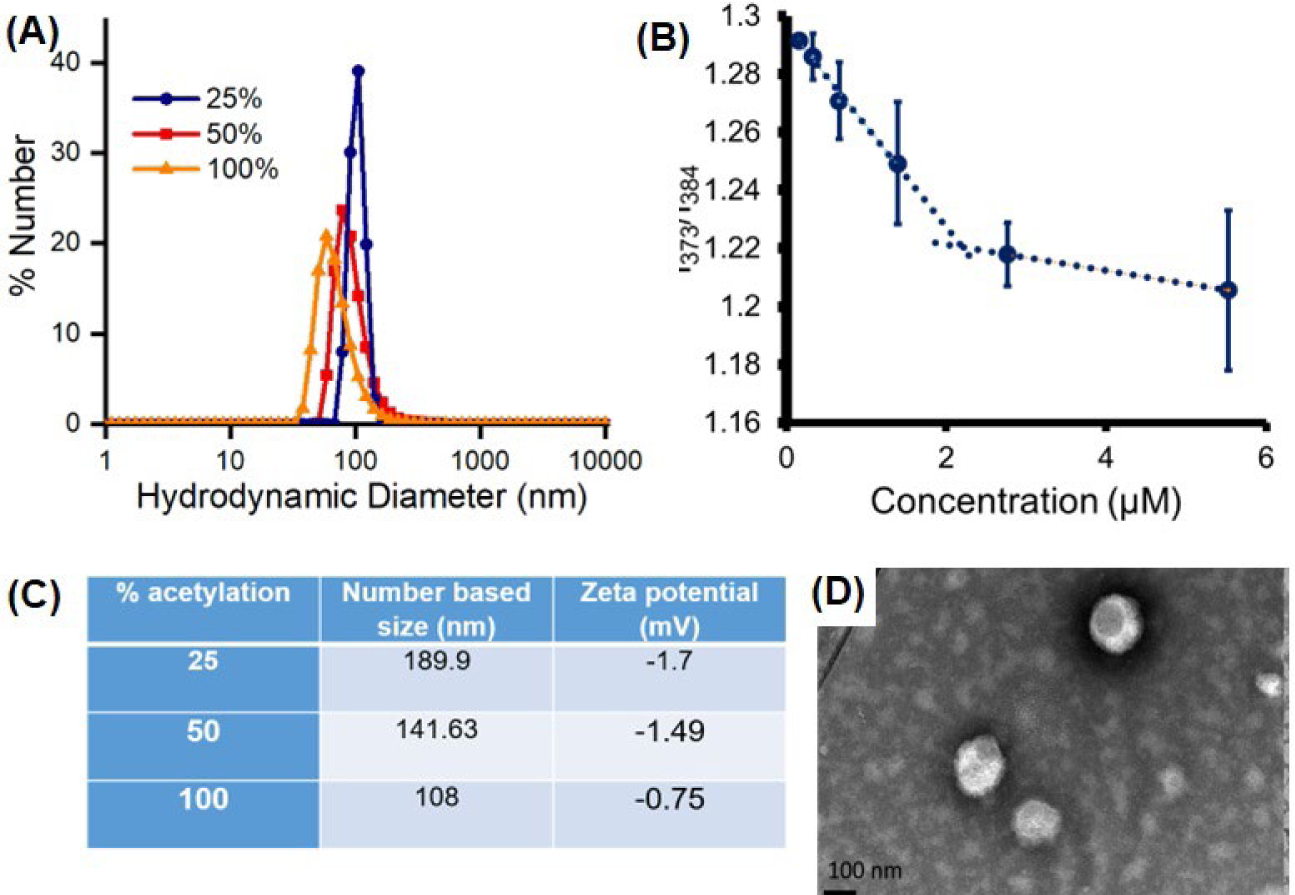
Nanoscale features of HDAC8-responsive particles (A) the hydrodynamic diameter of nanoparticles prepared from PEG-block-poly(acetylated L-lysine) as measured via DLS. Particle sizes were found to be governed by the degree of acetylation of the hydrophobic block. (B) Numerical values of the hydrodynamic diameters of HDAC8-responsive particles were calculated from the size distribution plot (A), along with surface charge (ζ-) potential values. (C) These particles are formed from the block copolymers aggregated in water above their critical aggregation concentration (CAC), calculated using pyrene-based fluorometric methods (D) Transmission electron microscopy (TEM) of HDAC8-responsive nanoparticles composed of PEG-block-poly (acetylated L-lysine).

We found that nanoparticles composed of PEG-block-poly (acetylated L-lysine) have a unimodal size distribution (**Figure 2A**). The degree of acetylation of the block copolymer influenced the mean hydrodynamic diameter of the nanoparticles formed. For example, nanoparticles composed of PEG-block-poly (acetylated L-lysine) at 25% and 50% degrees of acetylation resulted in the formation of nanoparticles with a hydrodynamic diameter of 190 and 142 nm, respectively. On the other hand, nanoparticles composed of PEG-block-poly (acetylated L-lysine) with 113 ethylene glycol units and 172 acetylated lysines (86% degree of functionalization) showed a hydrodynamic diameter of 108 ± 10.7 nm, with a polydispersity index (PDI) value of 0.248 (**Figure 2A**). The critical aggregation concentration (CAC) of these particles were found to be within the micromolar range (*2 x 10^-6^ M*, **Figure 2B**). Similarly, the ζ-potential measurement revealed that the nanoparticles formulated with copolymers with 25, 50 and 86% degree of acetylation showed ζ-potential of -1.7, -1.5 and -0.5 mV, respectively (**Figure 2C**). We observed that the ζ-potential of these systems is low, which could potentially impact the kinetic stability of the particles. However, we also noticed that the outer PEG corona of these particles imparts enough thermodynamic stability to the system, allowing them to maintain their stable colloidal structure for extended period of time (> 6 months). This morphology remained stable for an extended time, exceeding one year, under neutral pH conditions, as confirmed by TEM analysis. We used PEG-block-poly (acetylated L-lysine), with an 86% degree of functionalization, for transmission electron microscopy (TEM) and all other experiments. TEM experiments revealed that these block copolymers formed a distinct population of nanoparticles at pH 7.4 (TEM-based particle size = 100 nm, **Figure 2D**).

### 3.3. HDAC8 mediates Nanoparticle Destabilization

First, we set out to adjust the nanoparticle formulation for minimum background leakage of content and optimum sensitivity to HDAC8. For this reason, carboxyfluorescein-loaded nanoparticles composed of PEG-block-poly (acetylated L-lysine) were prepared. In 50 mM Tris-HCl buffer, containing 137 mM NaCl, 2.7 mM KCl, 1 mM MgCl2, and 1 mg/ml BSA maintained at pH 7.5 (later termed as HDAC-assay buffer), the nanoparticles released <5% of their contents (carboxyfluorescein) within 1h at 37°C in the absence of HDAC8 (**Figure 3A**). On the contrary, we observed that both the extent and rate of content release from the nanoparticles were increased with increasing HDAC8 concentration. We observed that 8, 15, and 45% of the loaded carboxyfluorescein was released when the nanoparticle formulations were incubated with 50 nm, 100 nm, and 1µM of HDAC8 under the same condition. We also observed that dye release from HDAC8-responsive nanoparticles can be significantly abrogated by introducing Suberoylanilide hydroxamic acid (SAHA), an HDAC8-specific inhibitor, in the release media at a concentration of 1µM along with HDAC8 (**Figure 3B**). Literature reports show that SAHA (also known as Vorinostat®) acts as an inhibitor of HDACs in both cytoplasm and nucleus.^[59]^ As a reversible and competitive inhibitor, SAHA binds to the zinc ion in the catalytic domain of HDACs, suppressing its enzymatic activity. Further, kinetic and thermodynamic studies reported earlier clearly showed that SAHA, compared to its structural analog Trichostatin A (TSA), binds preferentially to HDAC8.^[60]^ Concentration-dependence of CF release from the nanoparticles with HDAC8 and subsequent release inhibition upon incubation of the particles with SAHA provided us with an early indication regarding the responsivity of these nanoparticles towards the HDAC8 enzyme. This is because, in the presence of the inhibitor, there are few enzymes available to cleave off the acetyl groups from the poly (acetylated L-lysine) domain of nanoparticle-forming block copolymers, thereby exhibiting almost no increase in emission intensity of the dye over time.

**Figure 3.**
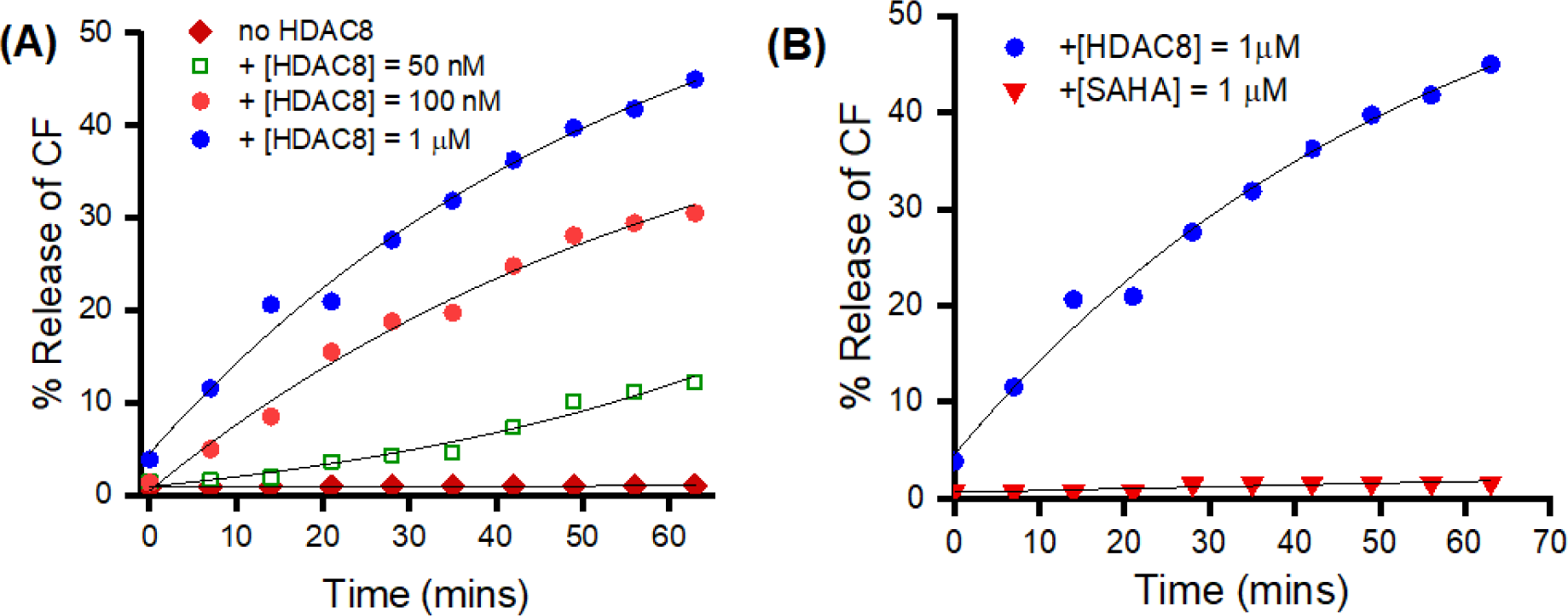
HDAC8-dependent release of encapsulated carboxyfluorescein (CF) from nanoparticles composed of PEG-block-poly(acetylated L-lysine). The kinetic trace of carboxyfluorescein fluorescence (*λ*_ex_ = 480 nm, *λ*_em_ = 518 nm) was monitored for 70 min for nanoparticles in the absence (green trace) and presence of a different concentration of HDAC8. The reactions were conducted at 25 °C in 50 mM Tris-HCl buffer containing 137 mM NaCl, 2.7 mM KCl, 1 mM MgCl_2_, and 1 mg/ml BSA maintained at pH 7.5. (B) Release of CF was abrogated when nanoparticles were co-incubated with SAHA (1µM), an HDAC8-specific inhibitor.

We further proved that when the nanoparticles were incubated with thermally deactivated HDAC8, no release of carboxyfluorescein was evident from within the particles. Nanoparticles loaded with carboxyfluorescein were treated with equimolar concentration of deactivated (heat-denatured) HDAC8 enzyme (**Figure 4A**). This evidence collectively indicates that nanoparticles formed from PEG-block-poly (acetylated L-lysine) favor releasing its encapsulated content only when incubated with HDAC8. Thus, their specific targeted ability to release encapsulated content in target cells.

**Figure 4.**
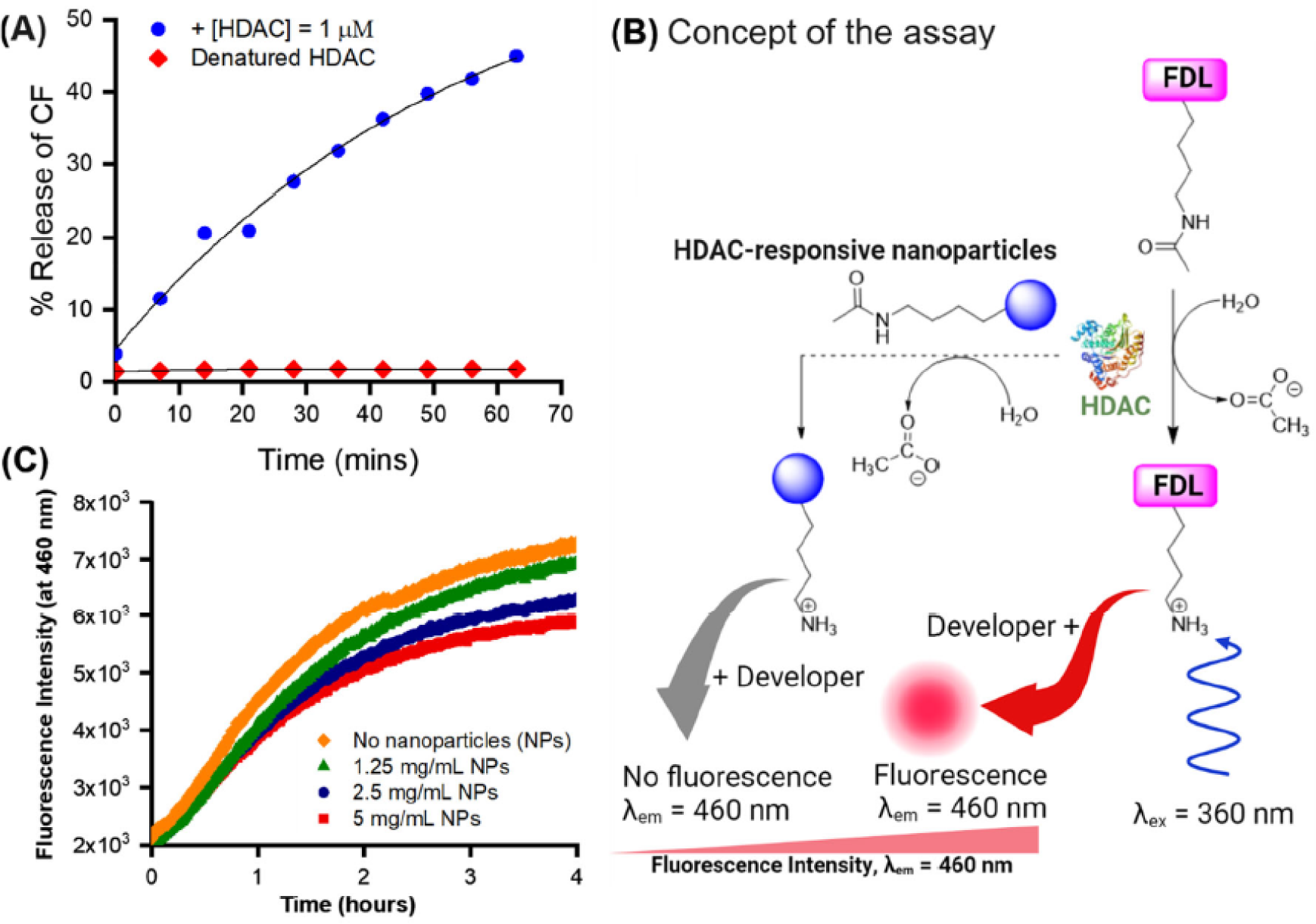
(A) Comparison of the release profile of carboxyfluorescein dye from nanoparticles triggered by HDAC8 (1µM) or by heat-denatured (deactivated) HDAC8 of equimolar concentration. (**B**) Mechanism of action of Fleur-de-Lys assay in the presence of HDAC enzymes. (**C**) Increasing the concentration of HDAC8-responsive nanoparticles decreases the fluorescence intensity to 460 nm, indicating the nanoparticles compete with the fluorogenic substrate for the enzyme.

We further set out to assess if the observed HDAC8-induced destabilization of nanoparticles is due to enzymatic interactions of the nanoparticles with HDAC8. We adopted a fluorometric assay using Fluor-de-Lys® (Enzo Life Sciences Inc.) as the fluorogenic substrate.^[16, 61]^ When HDAC8 binds to the reagent, fluorescence emission at 460 nm will increase due to the substrate’s cleavage from the reporter molecule. We used this assay to identify the binding of the nanoparticle with HDAC. The central concept of the assay is that, increasing the concentration of HDAC8-responsive nanoparticles decreases the fluorescence intensity to 460 nm, indicating the nanoparticles compete with the fluorogenic substrate for the enzyme (**Figure 4B**). In other words, the interaction between the nanoparticles and HDAC8 will reduce HDAC8 availability to the Fluor-de-Lys reagent, thereby reducing the fluorescence intensity of the reporter molecule. For this experiment, different concentrations of HDAC8-responsive nanoparticles were treated with HDAC8 in the presence of the Fluor-de-Lys substrate. We indeed observed that the emission intensity decreased with increasing particle concentration, indicating a possible competition of the enzyme-responsive nanoparticles with the reagent towards the enzyme (**Figure 4C**).

We investigated the destabilization properties of HDAC8-responsive nanoparticles under the influence of the enzyme using Atomic Force Microscopy (AFM) and nanoparticle tracking analysis (NTA). First, the AFM image of the untreated nanoparticles shows the typical spherical shape and dimension (**Figure 5A**). In contrast, the particle size was substantially decreased after treatment with 800 nM HDAC8 (**Figure 5B**), indicating HDAC8-driven destabilization of the particles. While the AFM images clearly show a change in size upon HDAC8 treatment, the NTA on the particle suspension provides the quantitative distribution of the particle size before and after treatments. From NTA measurements, we observed the particle concentration for untreated nanoparticles to be 1.7 × 10^7^ ± 2.12 × 10^6^ particles/mL, along with an average diameter of 160 ± 8 nm. After HDAC8 treatment for 1h, significant changes in the size and concentration of the particles were observed. The average particle size was measured to be 76 ± 33 nm, resulting in the shift of the particle distribution to the smaller fragment, most likely due to the dissociation of the deacetylated copolymers from the assembled systems (**Figure 5B**). Further, approximately a 9-fold increment in increased identical nanoparticle concentration (15.6 × 10^7^ ± 0.43 × 10^5^ particles/mL) after HDAC8 treatments additionally supports HDAC8-driven particle dissociation.

**Figure 5.**
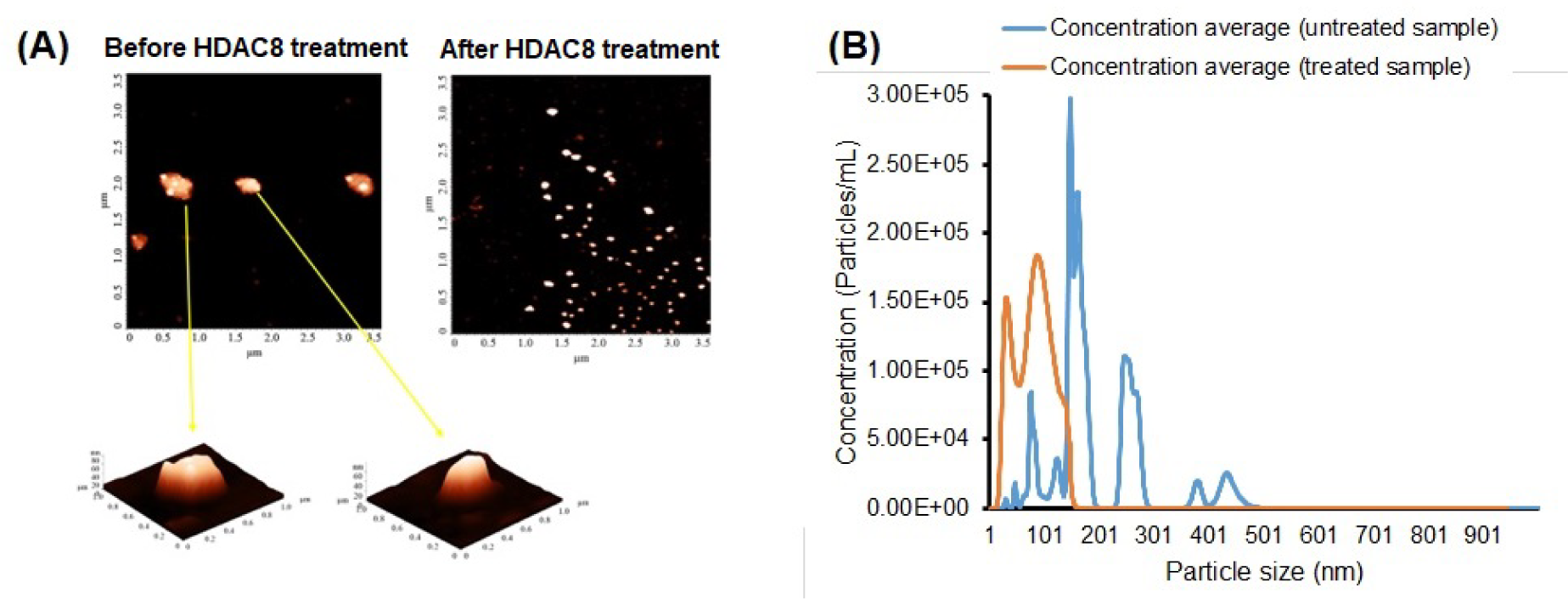
AFM topography of the nanoparticles and NTA analysis of the nanoparticle size distribution. (A) Nanoparticles before treating (left panel) and after treating with HDAC8 (800 nM) for 1h (right panel) showed a significant reduction in particle height and shape. (B) The average number and size of distribution of particles after HDAC8 treatments showed a reduction in particle size but a substantial increase in the concentration of reduced-size particles.

### 3.4. Proof-of-concept of using HDAC8-responsive nanoparticles as drug delivery platforms in cancer

Many types of cancers are HDAC-positive.^[21, 22]^ Most anticancer drugs present a narrow therapeutic window, and as such, targeted and triggered release of cytotoxic drugs to cancer tissue using nanoparticles is one of the successful and emerging therapeutic modalities against many types of cancer. Thus, using HDAC-responsive nanoparticles as platforms to encapsulate and release anticancer drugs seems viable to attain targeted therapy against HDAC-positive cancers.^[62-64]^ We set out to identify the potential of HDAC-responsive nanoparticles as PDAC-specific drug delivery. First, we identified the cytotoxicity of the polymer alone against cancer cell lines. We used three types of PDAC cells, namely, MIA PaCa-2, PANC-1, and pancreatic cancer stem cells (CSCs), to validate the proof-of-concept. We used non-neoplastic HPNE cells as a control. We showed that the acetylated block copolymer is not toxic to any of these cell lines, even at the micromolar concentration range **(Figure 6A)**. We identified a STAT3 inhibitor, Napabucasin (NAPA), as a model drug. The drug has been reported earlier to eliminate stemness-like tumor cells in different types of cancer. In addition, we encapsulated NAPA in HDAC-responsive nanoparticles. For this encapsulation method, both the copolymer, i.e., PEG-block-poly (acetylated L-lysine) and NAPA, were co-dissolved in DMSO, and solvent precipitated using buffer solution of pH 7.4 following our earlier published protocol.^[65-67]^ These drug-loaded nanoparticles were then assayed for their drug-loading and release efficiency triggered by HDAC8. We observed that HDAC8, the model enzyme, effectively triggered drug release from these NAPA-loaded nanoparticles within 4 h of incubation. In the absence of HDAC8, nanoparticles released < 10% of encapsulated NAPA, indicating the HDAC8-sensitive release of the encapsulated drug from these particles (**Figure 6B**). Our preliminary data showed that the drug-loaded nanoparticles showed a dose-dependent cytotoxicity against cancer cells and could exert a more potent cytotoxic effect on CSCs than other tested cell lines. We also noted the cytotoxic effect of NAPA on non-neoplastic cell lines, which could be attributed to the non-specific uptake of particles by these cells (**Figure 6C**). We are currently working on engineering more selectivity designed into these nanoparticles via adopting an active, ligand-mediated targeting approach.

**Figure 6.**
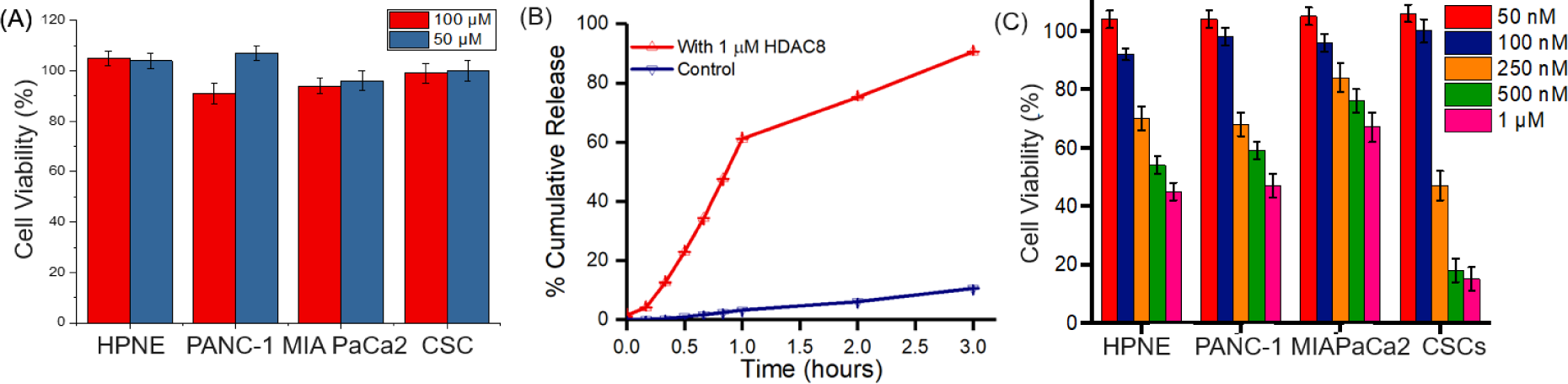
Interaction of HDAC8-responsive nanoparticles with different types of PDAC cells and non-cancerous HPNE cells in vitro. (A) HDAC8 (1 µM) triggers the release of NAPA, a selective STAT3 inhibitor from NAPA-encapsulated, HDAC8-responsive nanoparticles. Over 90% of the encapsulated drug is released from these nanoparticles after incubation with the enzyme for 3 h. Without the enzyme, the drug release rate and extent were significantly decreased. (B) NAPA-loaded nanoparticles showed a concentration-dependent effect on different types of cancer cells, with a more prominent effect on stem cells.

### 3.5. HDAC8 interactions in molecular docking studies

Based on the current structural design of block copolymers, exploring the structure-activity relationships of synthesized nanoparticles against human HDAC8 enzymes is of significant interest. We hypothesized that the nanoparticles interacted with the HDAC8 enzyme *via* the constituting block copolymers or unimers, i.e., PEG-block-poly (acetylated L-lysine). These unimers were docked into the active site of HDAC8 and compared with a native ligand, which corresponds to a sequence Arg379-His380-Lys381(ɛ-acetyl)-Lys382(ε-acetyl) of the p53 tumor suppressor protein.^[42]^ An X-ray crystal structure of HDAC8 in a complex with this chain (PDB ID: 2V5W) was selected for docking simulations. However, the target was reported with a Tyr306Phe mutation, which may affect the binding mode of the ligand. According to the previous investigations, the sidechain Tyr306 is crucial for the H-bonding interaction with acetyl moiety by adopting an ‘out-to-in’ transition in connection to a glycine-rich loop (Gly302-305), and any mutation in this HDAC8 loop is responsible to a genetic disorder, namely Cornelia de Lange Syndrome (CdLS).^[45, 68]^ Therefore, we rebuilt the HDAC8 model with Tyr306 by inducing a Phe306Tyr mutation in 2V5W, called Tyr306-2V5W. As a result, the protein structure after the mutation still highly overlapped with non-mutated HDAC8 (1T64) despite some conformational change (**Supporting information, Figure S3-S4**). While Phe152, Pro273, and Tyr306 residues remained in similar orientations, Asp101 displayed a large conformational change around the hinge region. The results suggested that Tyr306-2V5W was applicable for docking studies of PEG-b-p(LysAc) and the sidechain Asp101, as mentioned previously,^[42]^ could affect the binding mode of HDAC8 inhibitor.

To validate our docking approach, the co-crystal substrate was redocked into the active site of HDAC8 (**Figure 7A**). Please note that the co-crystal peptide was considered the native ligand of HDAC8 Tyr306-2V5W as there are similar structures before and after inducing Phe306Tyr substitution. After redocking the native ligand, the redocked and co-crystal conformers were highly superposed with a deviation value RMSD of 1.994Å. The fundamental interactions were conserved, including chelation with Zn^2+^, multiple H-bonds with Asp101 and Tyr306, and stacking interactions with Phe152, His180, and Phe208 (**Figure 7A**). The obtained results demonstrated the validity of the docking protocol established.

**Figure 7.**
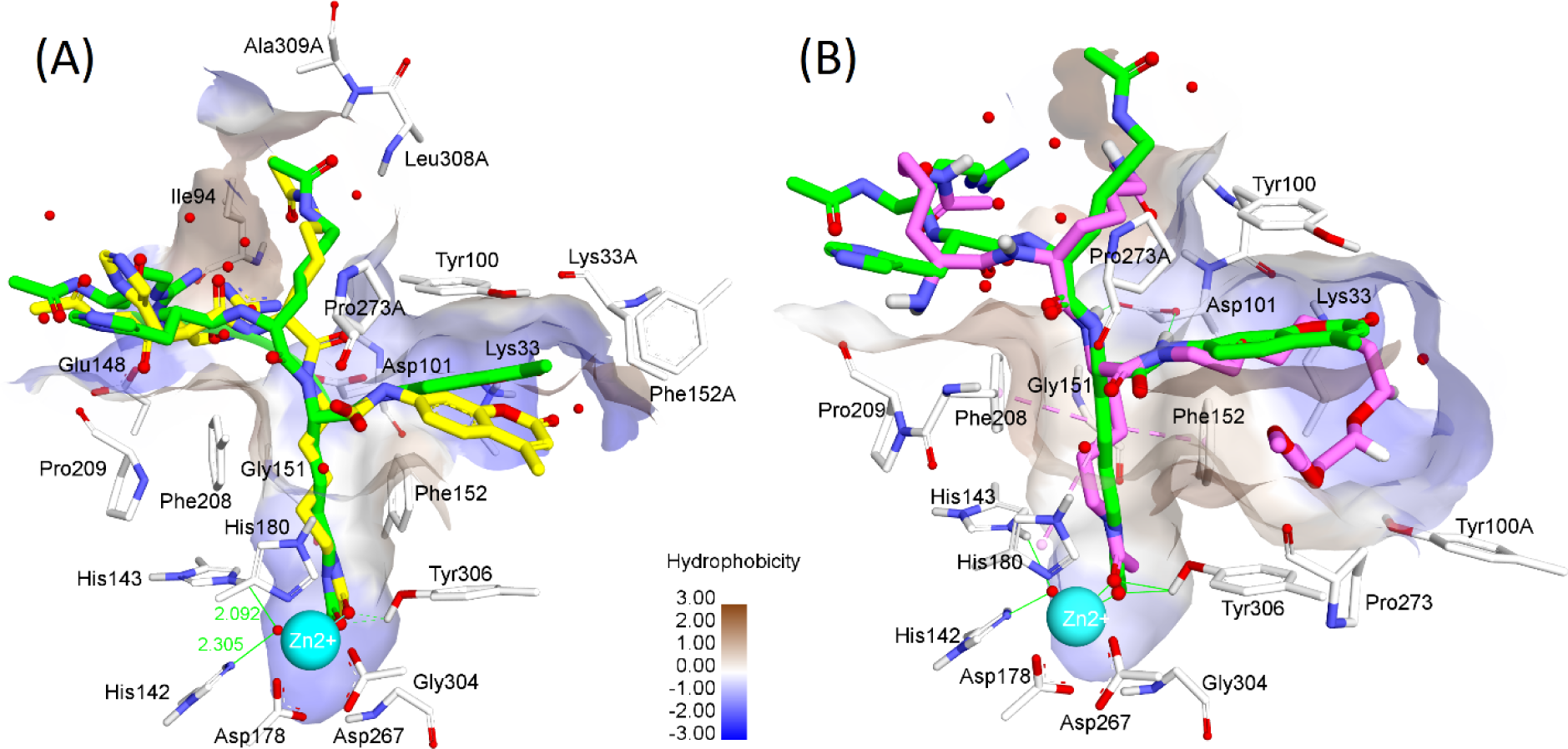
(A) superposition between redocked (yellow carbon) and co-crystal ligands (green carbon). (B) comparison between the binding modes of PEG-b-p(LysAc) (purple carbon) and co-crystal (green carbon) in the active site of HDAC8.

The next step involved docking PEG-b-p(LysAc) into the binding site of HDAC8 following the same protocol mentioned above (**Figure 7B**). To analyze the results, we divide the monomer into three parts according to the general HDAC inhibitor^[69]^ as illustrated in Figure 7 (i) Zn^2+^ binding group (ZBG) composed of N-acetyl group, (ii) the linker of 4C chain, and (iii) the capping group consists of PEG blocks in connection with two LysAc chains. At first sight, the docked monomer interacted with HDAC8 in a similar way as co-crystal peptides interacted. The N-acetyl lysine chain next to PEG accommodated well into the narrow cavity of the pocket and allowed the acyl group to chelate with Zn^2+^. The H-bond formed between the carbonyl oxygen of the acetyl group toward Tyr306 made the acyl group more susceptible to interacting with an activated water molecule that is Zn^2+^-coordinated and bound to His142 and His143 at the rim of the pocket. These interactions are crucial for the deacetylation reaction of HDAC8.^[45]^ According to the linker, stacking interactions with Phe152, His180, and Phe208 could keep the alkyl chain more stable in the tunnel.^[70]^ Several studies demonstrate the importance of hydrophobic interactions along the narrow tunnel of HDAC8 (**Figure 8**). The capping group, in turn, displayed diverse interactions. It would be remarked on the importance of Asp101 to form two H-bonds with two adjacent nitrogen atoms of the backbone Lysine. These interactions are crucial to keeping the ligand in the proper position in the pocket during the deacetylation reaction.^[42, 45]^ Of the capping group, two LysAc chains and PEG bound to different cavities (**Figure 8**). LysAc mainly showed interactions with Lys33, Tyr100, Pro273, etc. Meanwhile, the extended chain folding of PEG made the monomer able to interact with both hydrophobic and hydrophilic residues far from the pocket, including those of the other domains. The results suggest that, depending on the length of the PEG block (*x*) and the number of LysAc chains (*y*), the polymer could bind to a certain number of HDAC domains in a row, and the binding and releasing energies should be optimized based on (*x*, *y*) values. In addition, the affinities estimated by ICM scoring function for PEG-b-p(LysAc) and native ligand were quite similar, being -13.460 and -14.499 kcal/mol, respectively.

**Figure 8.**
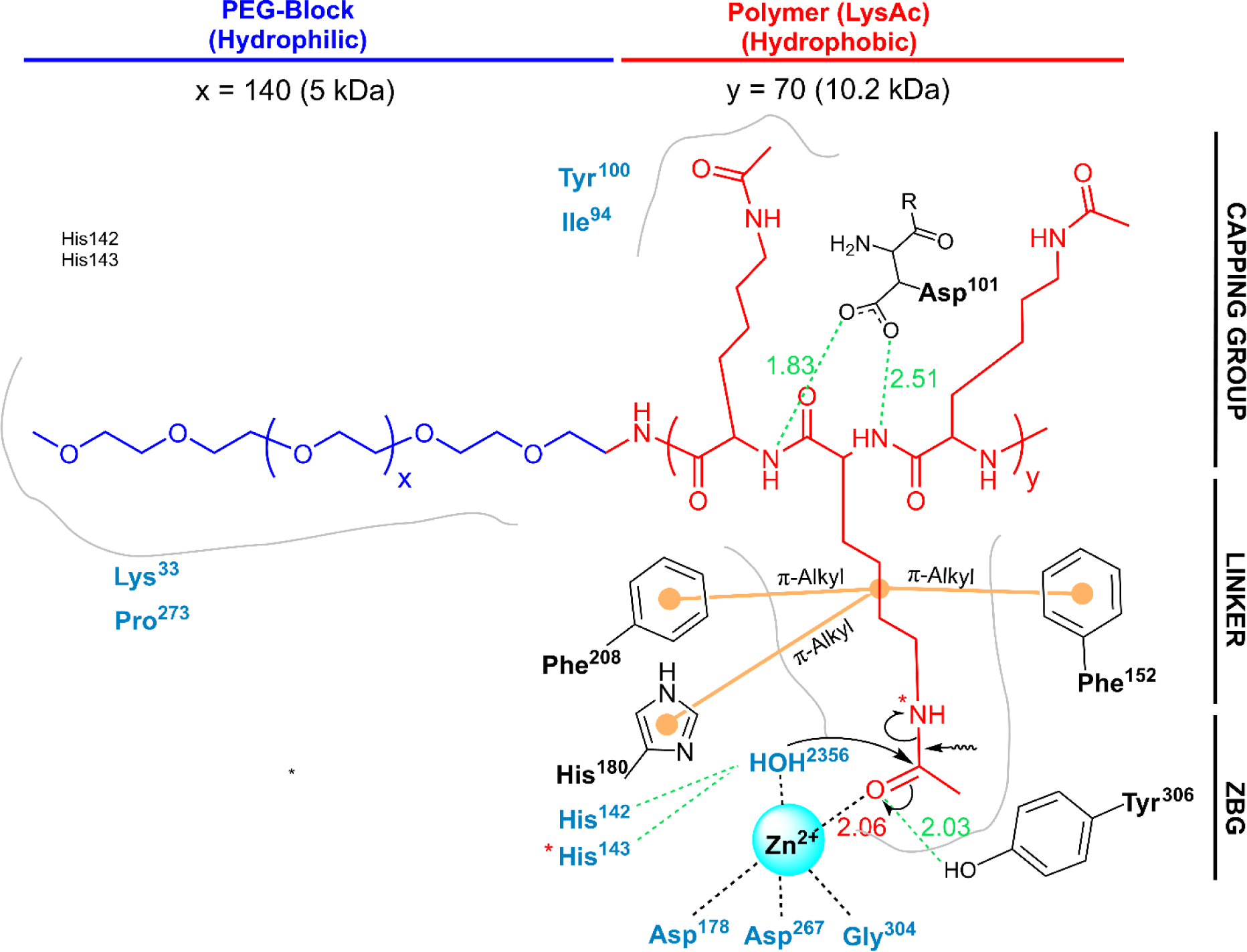
The critical interactions of PEG-b-p(LysAc) and residues in the binding site of HDAC8. Residues in black are crucial for binding. In green: H-bonds, orange: hydrophobic, black: electrostatic interactions. The mechanism of deacetylation is proposed according to Vannini *et al*.^[42]^

## 4. CONCLUSIONS

In summary, we demonstrated, for the first time, the design and development of HDAC-responsive nanoparticles, the contents of which can be released under the influence of a human HDAC enzyme. The nanoparticles were composed of PEG-block-poly(acetylated L-lysine) block copolymers. Due to a repeating sequence of acetylated L-lysine, the hydrophobic domain served as the HDAC-responsive site. As illustrated in this work using HDAC8 as the representative member of this enzyme family, the sensitivity of the nanoparticles towards HDAC8 was concentration-dependent and can be registered via the release of nanoparticle-encapsulated dye, carboxyfluorescein. We observed that ∼6 h was required to completely release the encapsulated content from the nanoparticle under the influence of HDAC8. In contrast, trypsin or serum albumin failed to initiate content release from the nanoparticles. This is most likely due to the deacetylation of the lysine residues from the hydrophobic domains of the nanoparticle-forming block copolymers. Such deacetylation can potentially lead to loss of amphiphilicity of the nanoparticle-forming block copolymer, creating irreparable defects within the particle structure that trigger the content release. As such, we envision that the HDAC-responsive nanoparticles could transport and release a drug payload to cellular targets, which overexpresses the HDAC family of enzymes. As a very early proof-of-concept, we showed that PDAC cancer-like stem cells, which overexpress HDACs, were more susceptible to anticancer drug therapy when delivered via HDAC-responsive nanoparticles. In addition, a computational molecular docking simulation confirmed the mechanism of interaction with the enzyme and strong binding affinity towards HDAC8. Overall, our methodology provides an early indication of finding a significant niche to target epigenetic enzymes, such as HDACs. Reminiscent of the naturally occurring histone protein, the nanoparticles synthesized in this work will pave the way for developing new functional materials that can be used to design artificial ‘histone-type’ organelles and epigenetics-based cellular networks. Currently, we are investigating the *in vivo* effect of HDAC8-responsive nanoparticles, which will bolster our findings to use HDAC-sensing particles in drug delivery, artificial organelle preparation, and diagnostic detection of genetically aberrant cell populations.

## ASSOCIATED CONTENT

### Supporting Information

The Supporting Information is available free of charge. NMR spectra, ninhydrin test protocols, sequence, and structural alignment of HDACs are available in supporting information.

### Author Contributions

The manuscript was written through the contributions of all authors. All authors have approved the final version of the manuscript.

### Funding Sources

This research was supported by NIH grant no. 2 P20 GM109024 from the National Institute of General Medicine (NIGMS), NSF CBET 2239626 (to M.Q.), NIH grant 2R01 GM114080 and NSF DMR 2322963 (to S.M.), NSF MCB 1942596 and CBET 2217474 (to Z.Y.). Partial support for this work was received from NSF grant no. IIA1355466 from the North Dakota Established Program to Stimulate Competitive Research (EPSCoR) through the Center for Sustainable Materials Science (CSMS). This work is also supported in part by the NSF MRI award OAC-2019077. Supercomputing support from the CCAST HPC System at NDSU is acknowledged. Funding for the Core Biology Facility used in this publication was made possible by NIH grant 2P20 RR015566 from the National Center for Research Resources. Its contents are solely the authors’ responsibility and do not necessarily represent the official views of the NIH. Any opinions, findings, conclusions, or recommendations expressed are those of the authors and do not necessarily reflect the views of the National Science Foundation.

### Notes

The authors declare no competing financial interest.

## ACKNOWLEDGMENTS

The authors thank Scott Payne and Jayma Moore of NDSU Electron Microscopy Core for TEM imaging.

## ABBREVIATIONS

HDAC8, histone deacetylase 8; TEM, transmission electron microscopy; DLS, dynamic light scattering; PEG, poly (ethylene glycol); CAC, critical aggregation concentration; PDAC, pancreatic ductal adenocarcinoma; CSC, cancer stem cells.

## Table of Content

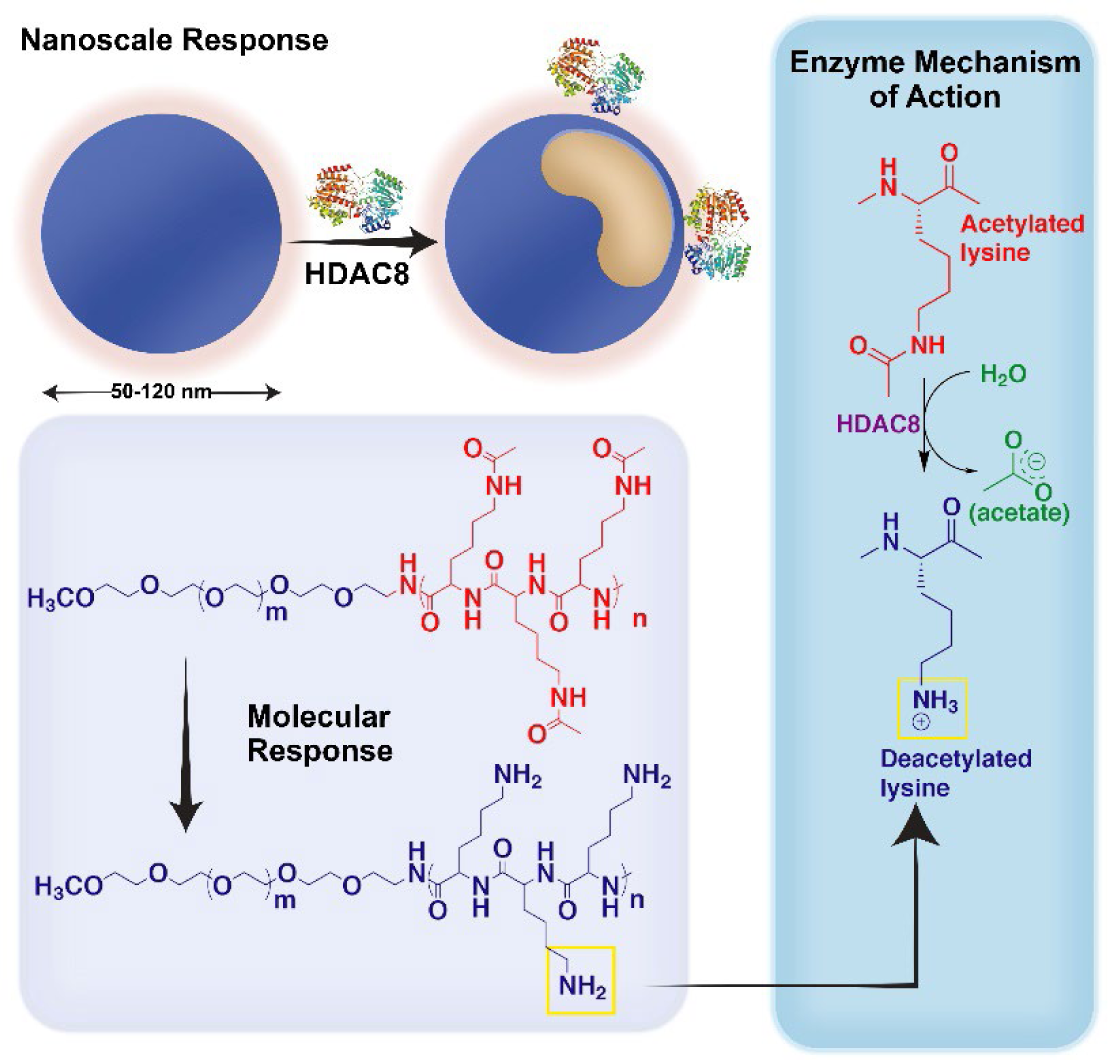

The work describes the synthesis of enzyme-responsive nanoparticles that respond to HDAC8, an enzyme of epigenetic class. The response mechanism is mediated by deacetylation reaction of acetylated Lysine, multivalently present within the hydrophobic domain of a block copolymer. First time report of these nanoparticles will open gateways to synthesize materials that can regulate epigenetic events and processes.

## REFERENCES

1. Song, S. J.; Choi, J. S., Pharmaceutics 2022, 14 (1), 143. DOI 10.3390/pharmaceutics14010143.

2. Yang, Y.; Aw, J.; Chen, K.; Liu, F.; Padmanabhan, P.; Hou, Y.; Cheng, Z.; Xing, B., Chem. Asian J. 2011, 6 (6), 1381–1389. DOI 10.1002/asia.201000905.

3. de la Rica, R.; Aili, D.; Stevens, M. M., Adv. Drug Deliv. Rev. 2012, 64 (11), 967–978. DOI 10.1016/j.addr.2012.01.002.

4. Rodriguez, A. R.; Kramer, J. R.; Deming, T. J., Biomacromolecules 2013, 14 (10), 3610–3614. DOI 10.1021/bm400971p.

5. Zhuang, J.; Seçinti, H.; Zhao, B.; Thayumanavan, S., Angew. Chem. Int. Ed. Engl. 2018, 57 (24), 7111–7115. DOI 10.1002/ange.201803029.

6. Moreno, S.; Sharan, P.; Engelke, J.; Gumz, H.; Boye, S.; Oertel, U.; Wang, P.; Banerjee, S.; Klajn, R.; Voit, B.; Lederer, A.; Appelhans, D., Small 2020, 16 (37), 2002135. DOI 10.1002/smll.202002135.

7. Wen, P.; Wang, X.; Moreno, S.; Boye, S.; Voigt, D.; Voit, B.; Huang, X.; Appelhans, D., Small 2021, 17 (7), 2005749. DOI 10.1002/smll.202005749.

8. Japir, A. A.-W. M. M.; Ke, W.; Li, J.; Mukerabigwi, J. F.; Ibrahim, A.; Wang, Y.; Li, X.; Zhou, Q.; Mohammed, F.; Ge, Z., J. Control Release 2021, 339, 418–429. DOI 10.1016/j.jconrel.2021.10.015.

9. Mura, S.; Nicolas, J.; Couvreur, P., Nature Mater. 2013, 12 (11), 991–1003. DOI 10.1038/nmat3776.

10. Hu, Q.; Sun, W.; Lu, Y.; Bomba, H. N.; Ye, Y.; Jiang, T.; Isaacson, A. J.; Gu, Z., Nano Lett. 2016, 16 (2), 1118–1126. DOI 10.1021/acs.nanolett.5b04343.

11. Randolph, L. M.; Chien, M.-P.; Gianneschi, N. C., Chem. Sci. 2012, 3 (5), 1363–1380. DOI 10.1039/C2SC00857B.

12. Kumar, V.; Munkhbat, O.; Secinti, H.; Thayumanavan, S., Chem. Commun. 2020, 56 (60), 8456–8459. DOI 10.1039/D0CC03257C.

13. Johnson, B. J.; Russ Algar, W.; Malanoski, A. P.; Ancona, M. G.; Medintz, I. L., Nano Today 2014, 9 (1), 102–131. DOI 10.1016/j.nantod.2014.02.005.

14. Paruchuri, B. C.; Gopal, V.; Sarupria, S.; Larsen, J., Nanomedicine 2021, 16 (30), 2679–2693. DOI 10.2217/nnm-2021-0194.

15. Wolffe, A. P., Science 1996, 272 (5260), 371–371. DOI 10.1126/science.272.5260.371.

16. Schultz, B. E.; Misialek, S.; Wu, J.; Tang, J.; Conn, M. T.; Tahilramani, R.; Wong, L., Biochemistry 2004, 43 (34), 11083–11091. DOI 10.1021/bi0494471.

17. Gregoretti, I.; Lee, Y.-M.; Goodson, H. V., J. Mol. Biol. 2004, 338 (1), 17–31. DOI 10.1016/j.jmb.2004.02.006.

18. Shahbazian, M. D.; Grunstein, M., Annu. Rev. Biochem. 2007, 76 (1), 75–100. DOI 10.1146/annurev.biochem.76.052705.162114.

19. Gantt, S. L.; Gattis, S. G.; Fierke, C. A., Biochemistry 2006, 45 (19), 6170–6178. DOI 10.1021/bi060212u.

20. Wolfson, N. A.; Pitcairn, C. A.; Fierke, C. A., Biopolymers 2013, 99 (2), 112–126. DOI 10.1002/bip.22135.

21. Ellis, L.; Atadja, P. W.; Johnstone, R. W., Mol. Cancer Ther. 2009, 8 (6), 1409–1420. DOI 10.1158/1535-7163.MCT-08-0860.

22. Milazzo, G.; Mercatelli, D.; Di Muzio, G.; Triboli, L.; De Rosa, P.; Perini, G.; Giorgi, F. M. Histone Deacetylases (HDACs): Evolution, Specificity, Role in Transcriptional Complexes, and Pharmacological Actionability Genes [Online], 2020.

23. Delcuve, G. P.; Khan, D. H.; Davie, J. R., Clin. Epigenet. 2012, 4 (1), 5. DOI 10.1186/1868-7083-4-5.

24. Yan, W.; Liu, S.; Xu, E.; Zhang, J.; Zhang, Y.; Chen, X.; Chen, X., Oncogene 2013, 32 (5), 599–609. DOI 10.1038/onc.2012.81.

25. Feng, W.; Zhang, B.; Cai, D.; Zou, X., Cancer Lett. 2014, 347 (2), 183–190. DOI 10.1016/j.canlet.2014.02.012.

26. Sethi, A.; Joshi, K.; Sasikala, K.; Alvala, M., Molecular docking in modern drug discovery: Principles and recent applications. Drug discovery and development-new advances. 2019; Vol. 2, pp 1–21.

27. Amaro, R. E.; Baudry, J.; Chodera, J.; Demir, Ö.; McCammon, J. A.; Miao, Y.; Smith, J. C., Biophysical journal 2018, 114 (10), 2271–2278.

28. Turabekova, M. A.; Rasulev, B. F.; Dzhakhangirov, F. N.; Leszczynska, D.; Leszczynski, J., Eur J Med Chem 2010, 45 (9), 3885–94. DOI 10.1016/j.ejmech.2010.05.042.

29. Turabekova, M.; Rasulev, B.; Theodore, M.; Jackman, J.; Leszczynska, D.; Leszczynski, J., Nanoscale 2014, 6 (7), 3488–95. DOI 10.1039/c3nr05772k.

30. Ahmed, L.; Rasulev, B.; Turabekova, M.; Leszczynska, D.; Leszczynski, J., Org Biomol Chem 2013, 11 (35), 5798–808. DOI 10.1039/c3ob40878g.

31. Yilmaz, H.; Ahmed, L.; Rasulev, B.; Leszczynski, J., Journal of Nanoparticle Research 2016, 18 (5), 123.

32. De Ruyck, J.; Brysbaert, G.; Blossey, R.; Lensink, M. F.; Chemistry, A. a. A. i. B. a., Molecular docking as a popular tool in drug design, an in silico travel. In Advances and Applications in Bioinformatics and Chemistry, 2016; pp 1–11.

33. Ren, Y.; Li, S.; Zhu, R.; Wan, C.; Song, D.; Zhu, J.; Cai, G.; Long, S.; Kong, L.; Yu, W., J. Med. Chem. 2021, 64 (11), 7468–7482. DOI 10.1021/acs.jmedchem.1c00136.

34. Thoma, G.; Patton, J. T.; Magnani, J. L.; Ernst, B.; Öhrlein, R.; Duthaler, R. O., J. Am. Chem. Soc. 1999, 121 (25), 5919–5929. DOI 10.1021/ja984183p.

35. Dan, K.; Bose, N.; Ghosh, S., Chem Commun (Camb) 2011, 47 (46), 12491–3. DOI 10.1039/c1cc15663b.

36. Bevilacqua, M. P.; Huang, D. J.; Wall, B. D.; Lane, S. J.; Edwards Iii, C. K.; Hanson, J. A.; Benitez, D.; Solomkin, J. S.; Deming, T. J., Macromol. Biosci. 2017, 17 (10), 1600492. DOI 10.1002/mabi.201600492.

37. Sanson, C.; Schatz, C.; Le Meins, J.-F.; Brûlet, A.; Soum, A.; Lecommandoux, S., Langmuir 2010, 26 (4), 2751–2760. DOI 10.1021/la902786t.

38. Gindy, M. E.; Ji, S.; Hoye, T. R.; Panagiotopoulos, A. Z.; Prud’homme, R. K., Biomacromolecules 2008, 9 (10), 2705–2711. DOI 10.1021/bm8002013.

39. Lebleu, C.; Rodrigues, L.; Guigner, J.-M.; Brûlet, A.; Garanger, E.; Lecommandoux, S., Langmuir 2019, 35 (41), 13364–13374. DOI 10.1021/acs.langmuir.9b02264.

40. Elegbede, A. I.; Banerjee, J.; Hanson, A. J.; Tobwala, S.; Ganguli, B.; Wang, R.; Lu, X.; Srivastava, D. K.; Mallik, S., J Am Chem Soc 2008, 130 (32), 10633–42. DOI 10.1021/ja801548g.

41. Abagyan, R.; Totrov, M.; Kuznetsov, D., J. Comput. Chem. 1994, 15 (5), 488–506. DOI 10.1002/jcc.540150503.

42. Vannini, A.; Volpari, C.; Gallinari, P.; Jones, P.; Mattu, M.; Carfí, A.; De Francesco, R.; Steinkühler, C.; Di Marco, S., EMBO Rep. 2007, 8 (9), 879–884. DOI 10.1038/sj.embor.7401047.

43. Somoza, J. R.; Skene, R. J.; Katz, B. A.; Mol, C.; Ho, J. D.; Jennings, A. J.; Luong, C.; Arvai, A.; Buggy, J. J.; Chi, E.; Tang, J.; Sang, B.-C.; Verner, E.; Wynands, R.; Leahy, E. M.; Dougan, D. R.; Snell, G.; Navre, M.; Knuth, M. W.; Swanson, R. V.; McRee, D. E.; Tari, L. W., Structure 2004, 12 (7), 1325–1334. DOI 10.1016/j.str.2004.04.012.

44. Neves, M. A. C.; Totrov, M.; Abagyan, R., J. Comput. Aided Mol. Des. 2012, 26 (6), 675–686. DOI 10.1007/s10822-012-9547-0.

45. Banerjee, S.; Adhikari, N.; Amin, S. A.; Jha, T., European journal of medicinal chemistry 2019, 164, 214–240. DOI 10.1016/j.ejmech.2018.12.039.

46. Ahmed, F.; Pakunlu, R. I.; Brannan, A.; Bates, F.; Minko, T.; Discher, D. E., J. Control. Release 2006, 116 (2), 150–158. DOI 10.1016/j.jconrel.2006.07.012.

47. Kim, J. O.; Nukolova, N. V.; Oberoi, H. S.; Kabanov, A. V.; Bronich, T. K., Polym. Sci. Ser. A 2009, 51 (6), 708–718. DOI 10.1134/S0965545X09060169.

48. Oh, K. T.; Bronich, T. K.; Bromberg, L.; Hatton, T. A.; Kabanov, A. V., J. Control. Release 2006, 115 (1), 9–17. DOI 10.1016/j.jconrel.2006.06.030.

49. Karandish, F.; Froberg, J.; Borowicz, P.; Wilkinson, J. C.; Choi, Y.; Mallik, S., Colloids Surf. B Biointerfaces 2018, 163, 225–235. DOI 10.1016/j.colsurfb.2017.12.036.

50. Visvader, J. E.; Lindeman, G. J., Nat. Rev. Cancer 2008, 8 (10), 755–768. DOI 10.1038/nrc2499.

51. Bao, B.; Ahmad, A.; Azmi, A. S.; Ali, S.; Sarkar, F. H., Curr. Protoc. Pharmacol. 2013, 61 (1), 14.25.1-14.25.14. DOI 10.1002/0471141755.ph1425s61.

52. Cruz, F. A.; Fonseca, A. N.; Moura, V.; Simoes, S.; Moreira, N. J., Curr. Pharm. Des. 2017, 23 (43), 6563–6572. DOI 10.2174/1381612823666171115105252.

53. Witt, A. E.; Lee, C. W.; Lee, T. I.; Azzam, D. J.; Wang, B.; Caslini, C.; Petrocca, F.; Grosso, J.; Jones, M.; Cohick, E. B.; Gropper, A. B.; Wahlestedt, C.; Richardson, A. L.; Shiekhattar, R.; Young, R. A.; Ince, T. A., Oncogene 2017, 36 (12), 1707–1720. DOI 10.1038/onc.2016.337.

54. Lin, P.-C.; Hsieh, H.-Y.; Chu, P.-C.; Chen, C. S. Therapeutic Opportunities of Targeting Histone Deacetylase Isoforms to Eradicate Cancer Stem Cells Int. J. Mol. Sci. [Online], 2018.

55. Chao, M.-W.; Chu, P.-C.; Chuang, H.-C.; Shen, F.-H.; Chou, C.-C.; Hsu, E.-C.; Himmel, L. E.; Huang, H.-L.; Tu, H.-J.; Kulp, S. K.; Teng, C.-M.; Chen, C.-S., Oncotarget 2015, 7 (2), 1796–1807. DOI 10.18632/oncotarget.6427.

56. Holowka, E. P.; Pochan, D. J.; Deming, T. J., J. Am. Chem. Soc. 2005, 127 (35), 12423–12428. DOI 10.1021/ja053557t.

57. Yaroslavov, A. A.; Yaroslavova, E. G.; Rakhnyanskaya, A. A.; Menger, F. M.; Kabanov, V. A., Colloids Surf. B Biointerfaces 1999, 16 (1), 29–43. DOI 10.1016/S0927-7765(99)00059-4.

58. Katayose, S.; Kataoka, K., Bioconjugate Chem. 1997, 8 (5), 702–707. DOI 10.1021/bc9701306.

59. Grabarska, A.; Łuszczki, J. J.; Nowosadzka, E.; Gumbarewicz, E.; Jeleniewicz, W.; Dmoszyńska-Graniczka, M.; Kowalczuk, K.; Kupisz, K.; Polberg, K.; Stepulak, A., J. Cancer 2017, 8 (1), 19–28. DOI 10.7150/jca.16655.

60. Singh, R. K.; Lall, N.; Leedahl, T. S.; McGillivray, A.; Mandal, T.; Haldar, M.; Mallik, S.; Cook, G.; Srivastava, D. K., Biochemistry 2013, 52 (45), 8139–8149. DOI 10.1021/bi400740x.

61. Wolfson, N. A.; Pitcairn, C. A.; Sullivan, E. D.; Joseph, C. G.; Fierke, C. A., Anal. Biochem. 2014, 456, 61–69. DOI 10.1016/j.ab.2014.03.012.

62. Ji, M.; Lee, E. J.; Kim, K. B.; Kim, Y.; Sung, R.; Lee, S.-J.; Kim, D. S.; Park, S. M., Oncol. Rep. 2015, 33 (5), 2299–2308. DOI 10.3892/or.2015.3879.

63. Zhou, Q.; Agoston, A. T.; Atadja, P.; Nelson, W. G.; Davidson, N. E., Mol. Cancer Res. 2008, 6 (5), 873–883. DOI 10.1158/1541-7786.MCR-07-0330.

64. Deng, B.; Luo, Q.; Halim, A.; Liu, Q.; Zhang, B.; Song, G., DNA Cell Biol. 2019, 39 (2), 167–176. DOI 10.1089/dna.2019.4877.

65. Ray, P.; Ferraro, M.; Haag, R.; Quadir, M., Macromol. Biosci. 2019, 19 (7), 1900073. DOI 10.1002/mabi.201900073.

66. Ray, P.; Kale, N.; Quadir, M., Colloids Surf. B Biointerfaces 2021, 200, 111563. DOI 10.1016/j.colsurfb.2021.111563.

67. Ray, P.; Dutta, D.; Haque, I.; Nair, G.; Mohammed, J.; Parmer, M.; Kale, N.; Orr, M.; Jain, P.; Banerjee, S.; Reindl, K. M.; Mallik, S.; Kambhampati, S.; Banerjee, S. K.; Quadir, M., Mol. Pharmaceutics 2021, 18 (1), 87–100. DOI 10.1021/acs.molpharmaceut.0c00499.

68. Porter, N. J.; Christianson, N. H.; Decroos, C.; Christianson, D. W., Biochemistry 2016, 55 (48), 6718–6729. DOI 10.1021/acs.biochem.6b01014.

69. Zhang, M.; Ying, J. B.; Wang, S. S.; He, D.; Zhu, H.; Zhang, C.; Tang, L.; Lin, R.; Zhang, Y., J. Cell. Biochem. 2020, 121 (5-6), 3162–3172. DOI 10.1002/jcb.29583.

70. Tabackman, A. A.; Frankson, R.; Marsan, E. S.; Perry, K.; Cole, K. E., J. Struct. Biol. 2016, 195 (3), 373–378. DOI 10.1016/j.jsb.2016.06.023.

