## Supporting information for "Design and Evaluation of Nanoscale Materials with Programmed Responsivity towards Epigenetic Enzymes"

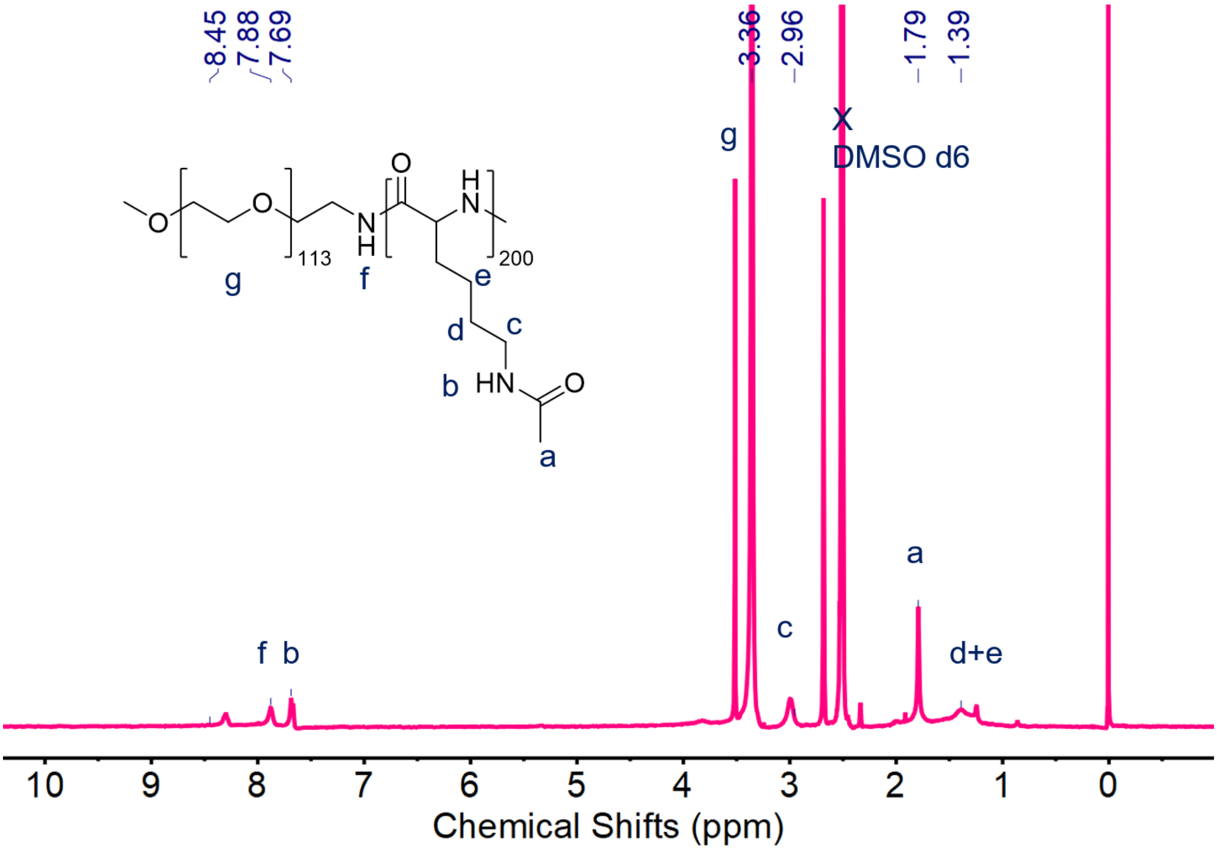


**Figure S1.** $^1$H NMR spectrum of acetylated PEG-b-poly(L-lysine) in DMSO-$d_6$)

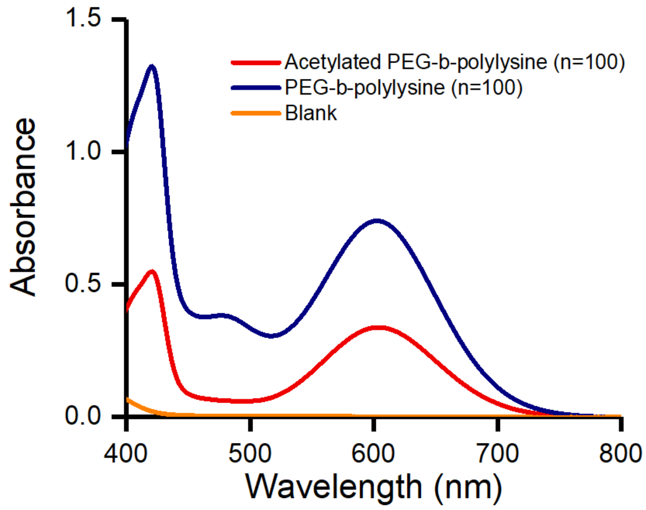


**Figure S2.** Absorbance spectra of block copolymers pre- and post-deacetylation using the ninhydrin test. The Ninhydrin test clearly shows the reduction of absorbance intensity originated from primary amines of PEG-block-poly(L-lysine) block copolymers due to acetylation. The Ninhydrin test was used to quantify the degree of functionalization in addition to $^1$H NMR.

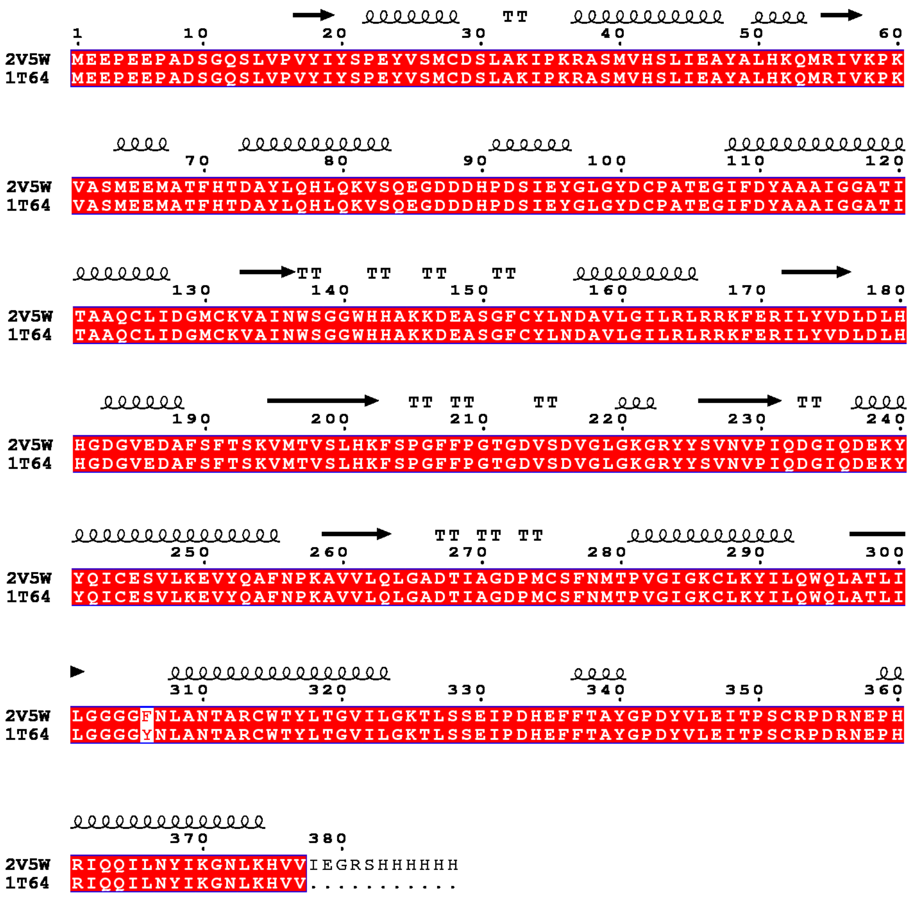


**Figure S3.** Sequence alignment between Tyr306Phe mutated (2V5W) and non-mutated (1T64) HDAC8.

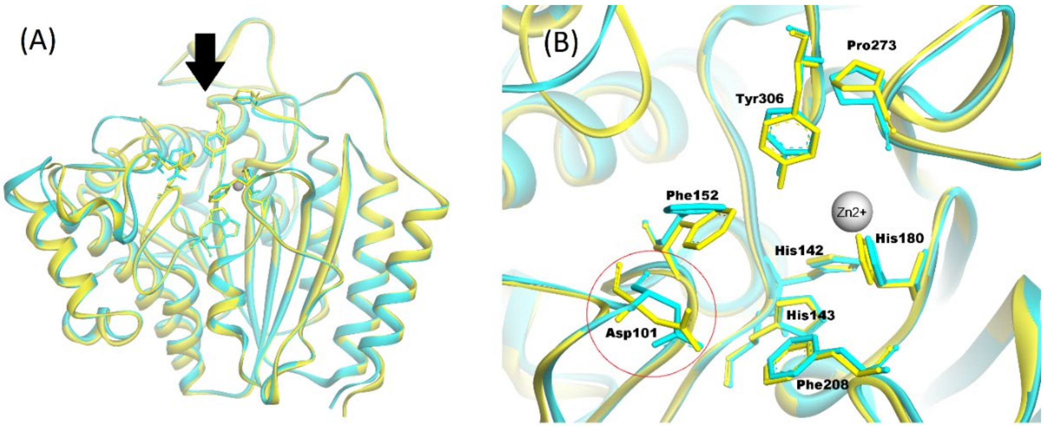

**Figure S4.** (A) Structural alignment of optimized Tyr$^{306}$-2V5W and non-mutated HDAC8 (1T64). (B) Visualization from the upper side (black arrow) revealed the movement of residues in the active site of two protein structures.
